# Targeting Heterochromatin Eliminates Chronic Myelomonocytic Leukemia Malignant Stem Cells Through Reactivation of Retroelements and Innate Immune pathways

**DOI:** 10.1101/2024.02.02.578583

**Authors:** Donia Hidaoui, Audrey Porquet, Rabie Chelbi, Mathieu Bohm, Aikaterini Polyzou, Vincent Alcazer, Stéphane Depil, Aygun Imanci, Margot Morabito, Aline Renneville, Dorothée Sélimoglu-Buet, Sylvain Thépot, Raphael Itzykson, Lucie Laplane, Nathalie Droin, Eirini Trompouki, Emilie Elvira-Matelot, Eric Solary, Françoise Porteu

**Affiliations:** Inserm UMR1287, Gustave Roussy Cancer Center, 94805, Villejuif, France; Université Paris-Saclay; Equipe labellisée Ligue Nationale Contre le Cancer; Inovarion, 75005 Paris, France; Institute for Research on Cancer and Aging of Nice (IRCAN), Université Côte d’Azur, INSERM, CNRS, Nice, France; Cancer Research Center of Lyon, UMR INSERM U1052 CNRS 5286, Centre Léon Bérard, Lyon, France; INSERM US23, CNRS UMS 3655, Gustave Roussy Cancer Center, Villejuif, France; Clinical Hematology Department, University Hospital, Angers, France; Hematology Department, Saint-Louis Hospital, Paris, France; Institut d’Histoire et Philosophie des Sciences et des Techniques, Université Paris I Panthéon-Sorbonne, Paris, France; Clinical Hematology Department, Gustave Roussy Cancer Center, Villejuif, France

## Abstract

Chronic myelomonocytic leukemia (CMML) is a severe myeloid malignancy affecting the elderly, for which therapeutic options are limited. DNA hypomethylating agents (HMAs) provide transient responses, failing to eradicate the malignant clone. Hematopoietic stem cell (HSC) aging involves heterochromatin reorganization, evidenced by alterations in histone marks H3K9me2 and H3K9me3. These repressive marks together with DNA methylation are essential for suppressing transposable elements (TEs). In solid cancers, the antitumor efficacy of HMAs involves the derepression of TEs, mimicking a state of viral infection. In this study, we demonstrate a significant disorganization of heterochromatin in CMML HSCs and progenitors (HSPCs) characterized by an increase in the repressive mark H3K9me2, mainly at the level of TEs, and a repression of immune and age-associated transcripts. Combining HMAs with G9A/GLP H3K9me2 methyltransferase inhibitors reactivates these pathways, selectively targeting mutated cells while preserving wild-type HSCs, thus offering new therapeutic avenues for this severe myeloid malignancy.

## Introduction

Chronic myelomonocytic leukemia (CMML) is a myeloid neoplasm originating from a hematopoietic stem cell (HSC). This disease usually emerges in the elderly (median age at diagnosis is 72 years) and combines dysplastic and proliferative features, therefore is classified as a myelodysplastic/myeloproliferative neoplasm^1^. Somatic mutations detected in CMML cells typically and diversely affect genes encoding epigenetic, splicing and signaling molecules. These mutations accumulate almost linearly in the stem cell compartment, with an early dominance of the clone that invades most of this compartment and a growth advantage to the most mutated cells with differentiation^2^. Such a relatively simple genotype contrasts with the heterogeneous phenotype of the disease. Consistently, abnormal DNA methylation profiles were identified as an additional feature of the disease^3–5^.

CMML is a severe neoplasm with a median survival lower than 3 years in most series. The sole curative therapeutic option is allogeneic stem cell transplantation, which is difficult to implement due to age and comorbidities. Cytoreductive drugs and DNA hypomethylating agents (HMAs) are used in patients with predominantly proliferative and dysplastic CMML variants, respectivey^6^. The therapeutic response to these drugs is always transient. The response to HMAs correlates with demethylation of leukemic cell DNA with a restored hematopoiesis that contrasts with the lack of decrease in the fraction of mutated cells^4^. Thus, HMA-induced correction of hematopoietic imbalance is likely attributable to epigenetic rather than cytotoxic effects.

In accordance with the age at diagnosis, two clock-like molecular signatures were identified in CMML cells^4^. Cellular aging also involves heterochromatin reorganization^7,8^, with alterations in DNA methylation and histone H3K9me2 and H3K9me3 mark patterns. Reduced H3K9me3 in HSCs is observed in aging or premature aging models, correlating with characteristic hematopoietic changes related to aging^9–11^. DNA methylation and repressive histone marks are crucial regulators of genome stability by repressing transposable elements (TEs), including DNA transposons and retrotransposable elements (RTEs). RTEs, which are classified into long terminal repeat (LTR) that characterize endogenous retroviruses (ERVs), and non-LTR elements such as long or short interspersed elements (LINE-1/L1; SINE), utilize a copy/paste mechanism to spread in the genome, causing double-strand breaks, and contribute to gene regulatory networks with a potential oncogenic role^12^. TEs also serve as a source of endogenous double-stranded (ds) RNA or cytoplasmic cDNA that are sensed as genomic parasites, ultimately triggering a type I interferon (IFN-I) response^13^. In HSCs, TE overexpression induced by irradiation and chemotherapy leads to DNA damage accumulation, and creates an intrinsic sterile inflammatory context potentially contributing to functional changes that generate an aging phenotype^14,15^.

HMA activity in solid tumors was ascribed to their capacity to derepress TEs that generate a type I interferon (IFN-I) and NFκB response and promote the expression tumor-associated antigens^16,17^. The role of these pathways in hematological malignancies treated with HMAs is less known. In the present study, we show a significant chromatin disorganization in HSPCs of CMML patients compared to aged healthy donors, which repress TE expression and inflammatory signaling. We also provide evidence that a combination of HMAs and H3K9me2 methyltranferase inhibitors could reactivate these pathways, specifically targeting the mutated clone while preserving unmutated HSCs.

## Results

### CMML HSPCs show a decrease in inflammatory response pathways

To explore the characteristics of pathological aged hematopoiesis in CMML patients versus healthy aging, we assembled a cohort of untreated CMML patients, at the time of diagnosis, and age-matched healthy donors (herein referred to as controls) (Supplementary Table 1). Bulk RNA-sequencing (RNA-seq) analysis was performed on CD34^+^ hematopoietic stem and progenitor cells (HSPCs) isolated from the bone marrow (BM) of 19 patients (mean age 70.6, range 49-84) and 12 age-matched healthy donors (mean age 68, range 54-91). Principal component analysis (PCA) showed separation of control and patient transcriptomes (Fig. 1a). Of 2,664 differentially expressed genes (DEGs) (adjusted pvalue <0.1), 54% were upregulated (Fig. 1b; supplementary Data 1). Gene set enrichment analysis (GSEA) examining HALLMARK and KEGG gene sets indicated that control cell transcriptomes were significantly enriched in various inflammatory, innate immune response, cytokine signaling, as well as cell cycle, P53, and apoptosis pathways (Fig. 1c; Supplementary Fig. 1a and Data 2). Gene Ontology process analysis unveiled that, beyond immune response, terms associated with chromatin remodeling and DNA packaging were enriched in controls (Fig. 1d; Supplementary Fig. 1b and Data 2). In contrast, patient cells exhibited enrichment in only a few terms related to ribosomes, vascular system development, actin, and adhesion (Fig. 1d; Supplementary Fig. 1a).

**Figure 1:**
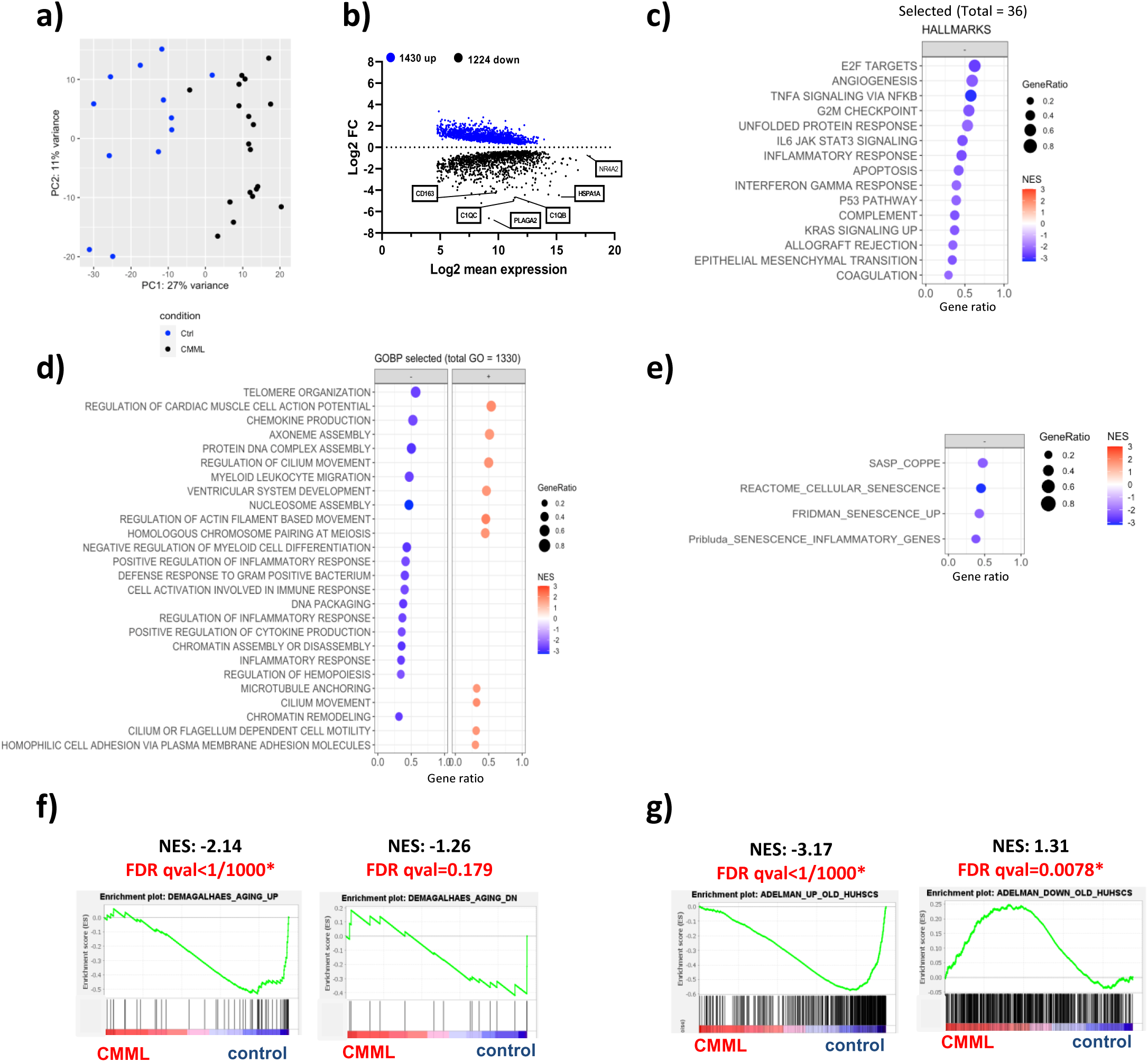
CMML HSPCs show a decrease in inflammatory response pathways. **a)** Principal component analysis (PCA) plot of BM CD34^+^ cells isolated from 19 CMML patients at diagnosis and 12 controls, based on RNA-seq regularized log transformed expression values. **b)** MA-plots showing differentially expressed genes (padj <0.1), upregulated (blue dots) or downregulated (black dots) in CMML cells. **c-e)** Dot plots showing selected enriched HALLMARK (total 36), GO (total 1330) and senescence gene sets upregulated (positive NES, red dots) and downregulated (negative NES, blue dots) in CMML cells. Gene sets are ranked by their NES and gene ration scores. **f, g)** Enrichment plots for aging published gene sets. FDR, false discover rate; NES, normalized enrichment score. * FDR <0.05

Increased expression of inflammatory genes, P53 signaling and chromatin deregulation are indicative of senescence and aging^8,18–22^.The transcriptome of patient HSPCs therefore suggests a younger state compared to control HSPCs. Supporting this possibility, senescence and aging signatures were more pronounced in control cells (Fig. 1e and f; Supplementary Fig. 1c). Furthermore, a signature composed of genes upregulated in aged versus young human BM HSCs^22^ was depleted in patient cells, while genes downregulated with age were marginally enriched (Fig. 1g).

### Chromatin is remodeled in CMML HSPCs

We have previously shown that murine and human HSC aging is associated with a loss of H3K9me3 heterochromatin mark^9,11^. Decreased expression of the heterochromatin mark H3K9me2, assessed by immunofluorescence, was also detected in HSCs from aged mice and in CD34^+^ BM cells from aged individuals compared with umbilical cord blood (Supplementary Fig. 2a and b). These results prompted us to analyze H3K9me3 and H3K9me2 expresssion in HSPCs from patients and controls. Surprisingly, immunofluorescence analysis suggested a significant decrease in H3K9me3 expression in patient cells (Fig. 2a), which was confirmed by immunoblot analysis in five patients (Supplementary Fig. 2c). As H3K9me3 is a repressive mark, this suggests that this epigenetic change is not responsible for the decreased expression of aging and senescence genes observed in cells from CMML patients, compared to age-matched control cells (Fig. 1f and g). In contrast, a significant increase in H3K9me2 expression was detected by immunofluorescence in the CD34^+^ cells of 28/29 patients (Fig. 2b), and validated by immunoblot analysis in five patients (Supplementary Fig. 2d). This correlated with heightened expression of the H3K9-dimethyltransferases G9A and GLP, particularly at the protein level (Fig. 2c and d; Supplementary 2e and f). Increased expression of H3K9me2, G9A and GLP were also observed in patient CD34^+^CD38^-^CD90^+^ HSC-enriched populations (Supplementary Fig. 2g-i). No correlation was found between the level of H3K9me2 in CD34^+^ cells and the presence of TET2, RAS, ASXL1 or SRSF2 mutations, WBC counts, the percent of BM blasts or age (Supplementary Fig2. j-m).

**Figure 2:**
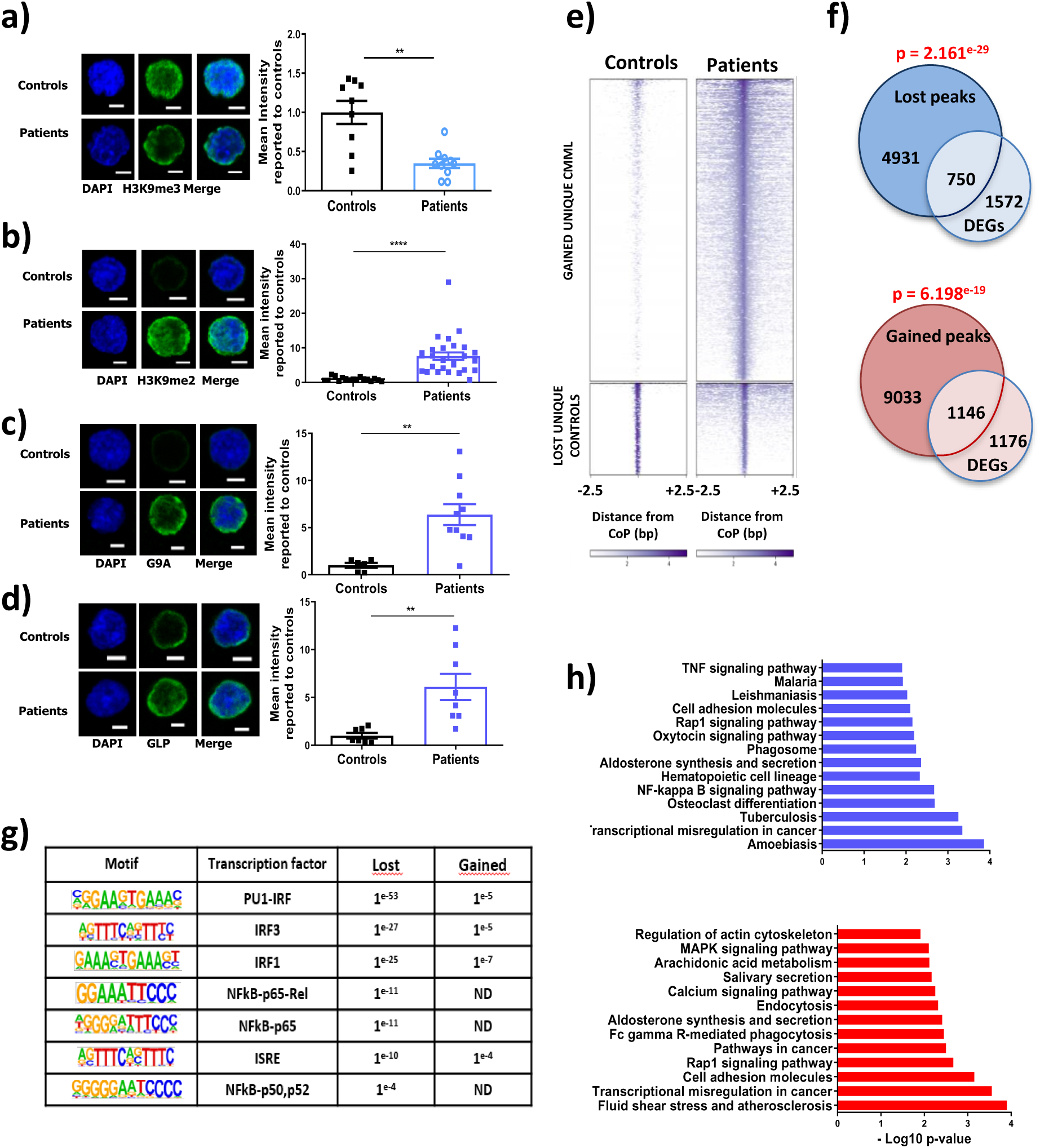
Chromatin is remodeled in CMML HSPCs. **a)** H3K9me3, **b)** H3K9me2, **c)** G9A and **d)** GLP mmunostaining in CD34^+^ cells from CMML patients and controls. Representative images and quantification of mean global immunofluorescence (IF) intensity using ImageJ software are shown. Means +/- SEM from 2-4 independent experiments normalized to the mean intensity of controls. Each dot represents cells from a single individual, with at least 50 cells counted per sample. Bars, 3 µm. Mann-Whitney test, **P<0.01; ****, P<0.0001. **e)** Heat map of the differentially chromatin accessible regions in CMML patient (n=4) and control (n=3) CD34^+^ cells +/- 2.5 kb from the center of the peak (CoP). **f)** Venn diagrams depicting the overlap of differentially expressed genes (DEGs) in CMML HSPCs with genes assigned to gained or lost ATAC peaks (-100/+25 kb from TSS). Fisher’s exact test. **g)** Motifs and p values for enrichment motifs of TF involed in IFN/NF-kB signaling in ATAC-seq peaks unique to patients (gained) and controls (lost) using HOMER. ND: non detected **h)** KEGG gene set analysis of deregulated genes that also exhibit gained (red) or loss chromatin accessibility in CMML patient cells (blue).

To explore the impact of these opposite modifications on two heterochromatin marks on chromatin opening genome-wide, we conducted Assay for Transposase-Accessible Chromatin-sequencing (ATAC-seq) experiments in CD34^+^ cells from 4 patients and 3 controls. We detected 40,335 differential peaks, with 31,110 (77%) gained and 9,245 (23%) lost in CMML cells (Fig. 2e; Supplementary Data 3). Altered peaks were significantly enriched in intergenic and intronic regions, while common peaks were predominantly located at the transcription start sites (Supplementary Fig. 3a). Assigning differential ATAC-seq peaks to genes^15^ demonstrated a substantial overlap between DEGs and ATAC-seq peaks (Fig. 2f). Analysis of transcription factor (TF) motifs in peaks using HOMER demonstrated specific or stronger enrichment of motifs for TF involved in hematopietic differentiation (FL11, RUNX1, GATA1, GATA1, NF-E2, EKLF, MAFB, MYB), TGF-β signaling (SMAD2, SMAD3, SMAD4) inflammation (NFκB, IRF1, IRF3, RFX, ISRE), in ATAC-seq peaks with reduced compared with gained chromatin accessibility (Fig. 2g; Supplementary Fig. 3b; Supplementary Data 4). This is consistent the lower enrichment of these pathways in CMML patient cells (Fig. 1c, d; Supplementary Data 2). In addition, numerous processes related to immune/inflammation/NFκB signaling were associated with genes exhibiting loss of chromatin accessibility (Fig. 2h; Supplementary Fig. 3c). This suggests that the attenuated expression of certain immune/inflammatory response genes in patient cells might be related to an altered chromatin accessibility.

Together, these results highlight a profound chromatin remodeling that may contribute to repressing inflammatory genes in CMML compared to healthy CD34^+^ cells.

### H3K9me2 is enriched at TEs in CMML HSPCs

H3K9me2 is recognized as a suppressor of innate immune genes, notably exerting direct control over the expression of *IFNB1* gene, encoding IFN-β, and certain IFN-stimulated genes (ISGs)^23,24^. This suggests that increased H3K9me2 in CMML cells could contribute to the observed decreased chromatin accessibility at immune/inflammatory genes and in their expression. We therefore focused on this mark, performing H3K9me2 CUT&Tag analysis in CD34^+^ cells, comparing 4 patients with 3 controls. As expected from immunofluorescence and western blot experiments, patient cells exhibited a greater number of recovered peaks (15,406 ± 1,484) compared to control cells (4,831 ± 1,600) (Supplementary Fig. 4a). Focusing on peaks shared by all four patients or three controls, we identified 1,396 gained peaks exclusive to patients and only 88 peaks unique to controls (Fig. 3a; Supplementary Data 5). Gene annotation revealed that H3K9me2 peaks spread mainly across intergenic and intronic regions in both patient and control cells (Fig. 3b). However, when compared to the genome using Genome Association Test, only a small significant enrichment of the mark at promoters and 1-5kb regions in patient cells (Supplementary Fig. 4b). Biological processes associated with the nearest promoters gaining H3K9me2 in patient cells did not show enrichment in immune pathway genes or IRF/NFκB transcription factors (Suppmentary Data 6). Moreover, there was no significant difference between patients and controls in the expression of H3K9me2-linked genes (Supplementary Fig. 4c), and no correlation was identified between H3K9me2 concentration at the promoter level and gene expression (Supplementary Fig. 4d). Finally, there was no enrichment of H3K9me2 reads at peaks showing either decreased or increased accessibility (supplementary Fig. 4e), suggesting that H3K9me2 increase in CMML patients is not directly linked to chromatin accessibility and gene expression.

**Figure 3:**
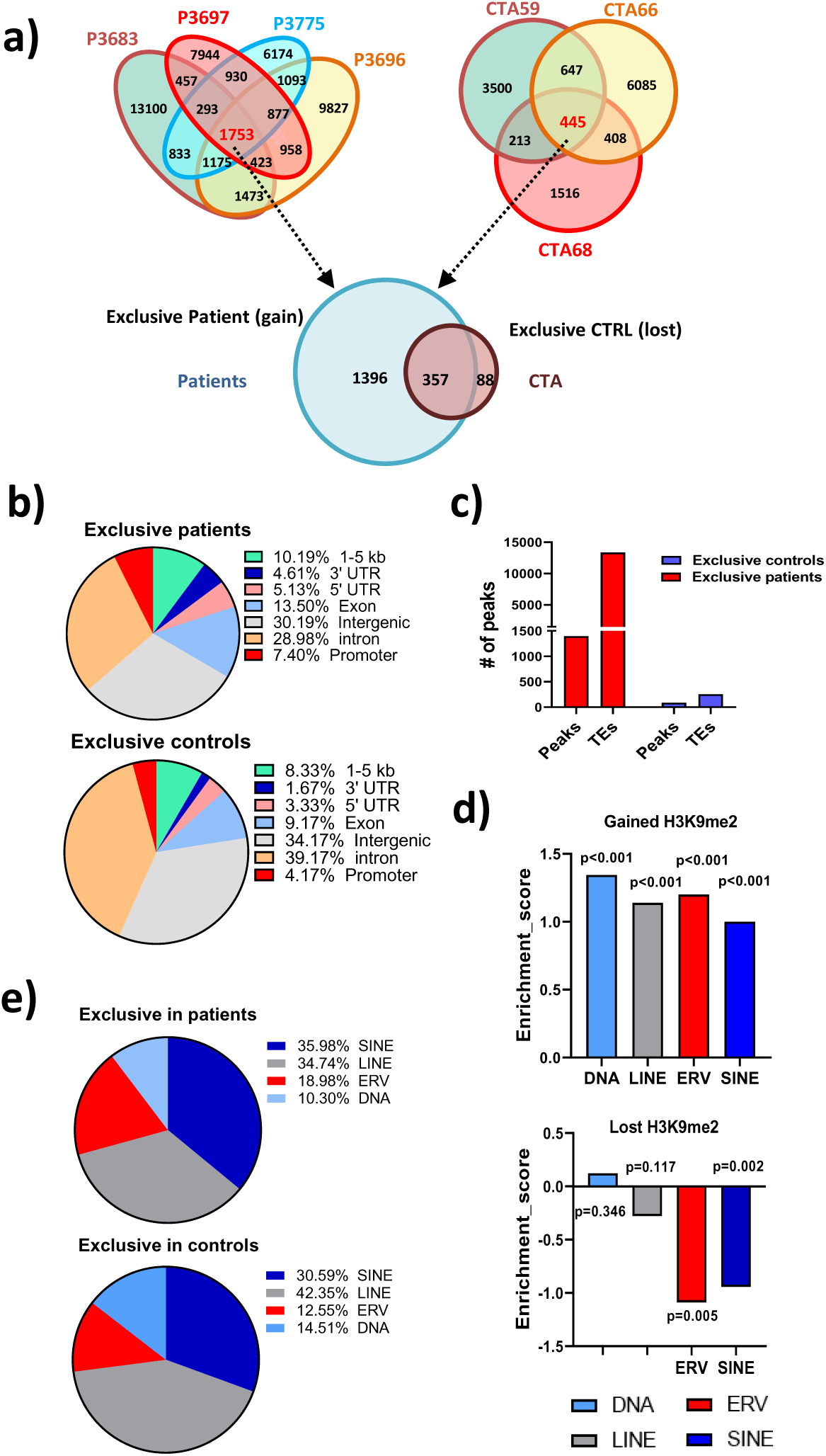
H3K9me2 is increased at TEs in CMML CD34^+^ cells. **a)** Venn diagram of peak retrieved from the calling from H3K9me2 CUT&Tag on CD34^+^ from 4 patients (P) and 3 controls (CTA). **b)** Repartition of the CUT&Tag peaks found exclusively in patients and controls. **c)** Numbers of common peaks found exclusively in the 4 patients (red) and 3 controls (blue) and numbers of TEs annotated in these peaks. **d)** Enrichment or depletion of the different classes of TEs in gained or lost H3K9me2-CUT&Tag peaks. The Y-axis representing the enrichment score was calculated as the log2 fold change of the number of specific peaks overlapping with TEs found in patient gained or lost peaks over the median number of TEs found in 1000 times the same number of randomly selected peaks overlapping with TEs. Positive log2 fold change = enrichment, and negative log2 fold change = depletion. **e)** Repartition of the different TE classes in CUT&Tag peaks found exclusively in patients and controls.

Since H3K9me2 is also known to repress TEs^25,26^, we analyzed CUT&Tag data by considering all mapping reads, either unique and multiple, that mapped without mismatch, and we randomly assigned them at one of their best possible positions in the genome, as described^11^. This analysis identified a pronounced enrichment of H3K9me2 at TEs relative to genes in patients, with 13,364 TEs among H3K9me2-bound peaks in CMML cells, compared to 255 in control cells (Fig. 3c; Supplementary Fig. 4f). Enrichment of TEs in patient and control peaks was performed using a permutation test, counting peaks that intersected with any TE belonging to a given TE class in CMML unique (gained) and control unique (lost) peaks and in 1000 times an equal number of peaks randomly chosen. This revealed that H3K9me2 peaks gained in patient cells were significantly enriched for all TE classes (Fig. 3d; Supplementary 4g). In contrast, ERVs and SINEs were under-represented in lost peaks. The distribution of H3K9me2 among TE classes showed that the proportion of ERVs was greater in peaks gained than in peaks lost, while LINEs were overrepresented among TE lost (Fig. 3e). Overall, these results highlight that the profound chromatin remodeling detected in CMML CD34^+^ cells involves an enrichment of H3K9me2 at TEs.

### The HMA and G9A/GLP inhibitor combination reduces CMML HSPC clonogenicity

Recent data have shown that HMAs, when used in various pathological contexts, can act through the reactivation of TEs^16,17^, ultimately triggering the activation of IFN-I and NF-κB pathways. Our initial results led us to speculate that deregulated G9A/GLP methyltransferases and histone marks could interfere with CMML cell response to HMAs. To test this hypothesis, CMML and healthy donor CD34^+^ cells were treated *ex vivo* with the G9A/GLP inhibitor UNC0638, the HMA decitabine (DAC) and their combination, then seeded in methylcellulose for 14 days (Fig. 4a). Since H3K9me2 has been shown to bind to demethylated regions where it can then reinstall DNA methylation^26,27^, we added UNC0638 2 days after DAC. Dose-response experiments demonstrated that each compound alone had a modest impact on colony formation by patient cells, with 33% and 6% decrease observed at the highest dose of DAC (80 nM) and UNC0638 (2 µM), respectively. In contrast, the combination of DAC and UNC0638, even at low doses, significantly reduced the clonogenicity of these cells (Fig. 4b). Using the Chou-Talalay method^28^, a synergistic effect was observed under all conditions (Fig. 4c and d). Consistent with the low expression of G9A/GLP in these cells, no synergy between DAC and UNC0638 on their clonogenicity was detected (Supplementary Fig. 5a). Although the highest dose of DAC showed some toxicity on healthy cells, CMML leukemic cells were much more sensitive to the DAC/UNC0638 combination, with the highest doses decreasing by 80% and 20% the number of colonies formed by CD34+ cells from patients and control cells, respectively.

**Figure 4:**
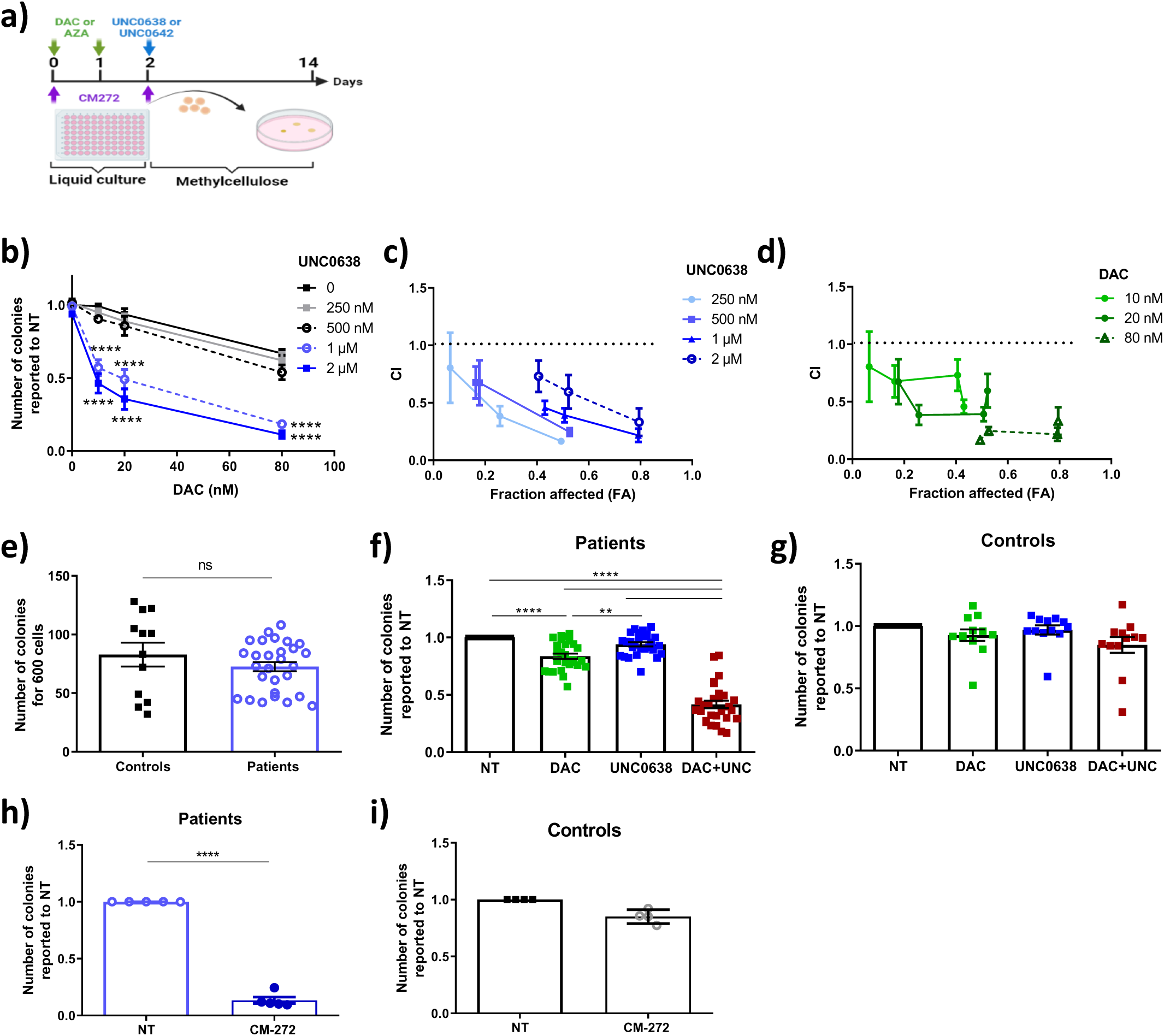
the HMA and G9A/GLP inhibitor combination reduces CMML HSPC clonogenicity. **a)** Treatment protocol of CD34^+^ cells. DAC: decitabine; AZA: 5-azacytidine; UNC0638 and UNC0642: G9A/GLP inhibitors; CM-272: dual DNMT1 and G9A/GLP inhibitor. **b)** Dose-response effect of DAC (X-axis) and UNC0638 (color scale) on CMML CD34^+^ colony formation. Mean +/- SEM from 3 patients. Two-way ANOVA Bonferroni’s multiple comparisons **c-d**) Chou-Talalay model of the effects of combining different doses of DAC and UNC0638. The fraction affected (fa) corresponds to the proportion of colonies eliminated by the treatments. The combination index (CI) is calculated by CompuSyn software (ComboSyn. Inc.) where CI<1, CI=1, CI>1 indicates a synergistic, additive and antagonistic effect respectively. Mean+/- SEM from 3 patients. e) Number of colonies formed from CMML (n=28) and control (n=12) CD34^+^ cells in the absence of treatment. One dot represents a single individual. t-test **f, g)** Number of colonies formed by CD34^+^ cells from patients (f) or controls (g) after treatment with 10 nM DAC and 1μM UNC0638 alone or in combination. NT: non-treated. One dot represents a single individual; controls, n=12; patients, n=28. Results are normalized to the number of colonies in the NT condition. Means +/- SEM. One-way ANOVA Bonferroni’s Multiple Comparison. **h, i)** Number of colonies formed by CD34^+^ cells from patients, n=5 (h) or controls, n=4, (i) after treatment with or without (NT) 200 nM CM-272. Results are normalized to the number of colonies in the NT condition. One dot represents a single individual. Means +/- SEM. t.test. **p<0.01; ***p<0.001; **** p<0.0001.

Subsequent experiments were performed with 10 nM DAC followed by 1 µM UNC0638. A significant reduction in global DNA methylation level was detected by ELISA after 2 days of DAC, which was maintained at day 4, either alone or combined with UNC0638, while UNC0638 alone had no effect on DNA methylation (Supplementary Fig. 5b and c). The level of H3K9me2, measured by immunofluorescence, remained unchanged after 2 days of culture in the presence of DAC alone (Supplementary Fig. 5d), while it was markedly reduced by UNC0638 (Supplementary Fig. 5e). These experiments validated the proper functioning of the tested drugs.

In the absence of treatment, the numbers of colonies formed by CD34^+^ cells were comparable between control and patient cells (Fig. 4e). Treatment with DAC+UNC0638 inhibited the clonogenicity of 26 out of 28 tested CMML CD34^+^ cells and reduced the colony size while each inhibitor alone had a weaker (DAC) or no (UNC0638) effect (Fig. 4f; Supplementary Fig. 6a and b). Again, none of these treatments affected the growth of control CD34^+^ cells (Fig. 4g, Supplementary Fig. 6a). Similar results were obtained with the combination of another HMA, 5-Azacytidine (AZA) combined with UNC0638, and with another G9A/GLP inhibitor, UNC0642, combined with DAC (Supplementary Fig. 6c-f). Finally, CM-272, a dual inhibitor of DNMT1 and G9A/GLP^29^, specifically reduced the clonogenicity of CMML but not control CD34^+^ cells (Fig. 4h and i).

To more precisely identify the cell population targeted by the combination, CD34^+^ cells were enriched in CD34^+^CD38^-^CD90^+^ (HSC-enriched, referred to as stem cells) and CD34^+^CD38^-^CD90^-^ (progenitor-enriched, referred to as progenitors) cells. We functionally assessed how the inhibitors affected the cell clonogenic potential by testing their capacity to serially replate in methylcellulose over 3 passages after treatment at day 2 and 4 of the culture (Fig. 5a). In the absence of treatment, at all three passages, the number of colonies formed by control and patient stem cells was not significantly different (Supplementary Fig. 7a and b). The DAC/UNC0638 combination triggered a drastic reduction in the number and size of colonies formed by both CMML stem cells and CMML progenitors (Fig. 5b and Fupplementary Fig. 7b and c). DAC alone slightly decreased the size of the colonies generated by CMML progenitors (Supplementary 7b) and, starting from P2, significantly reduced their number, while having no effect on the stem cell-enriched population (Fig. 5b). Measuring the cumulative clonogenic potential at P2, which considers both the number of colonies and their size^30^, confirmed the strong effect of the combination on both stem cells and progenitors while DAC affects in priority the latter populations (Fig. 5c). None of the treatments affected colony formation by stem cells or progenitors from controls (Supplementary Fig. 7c and d).

**Figure 5:**
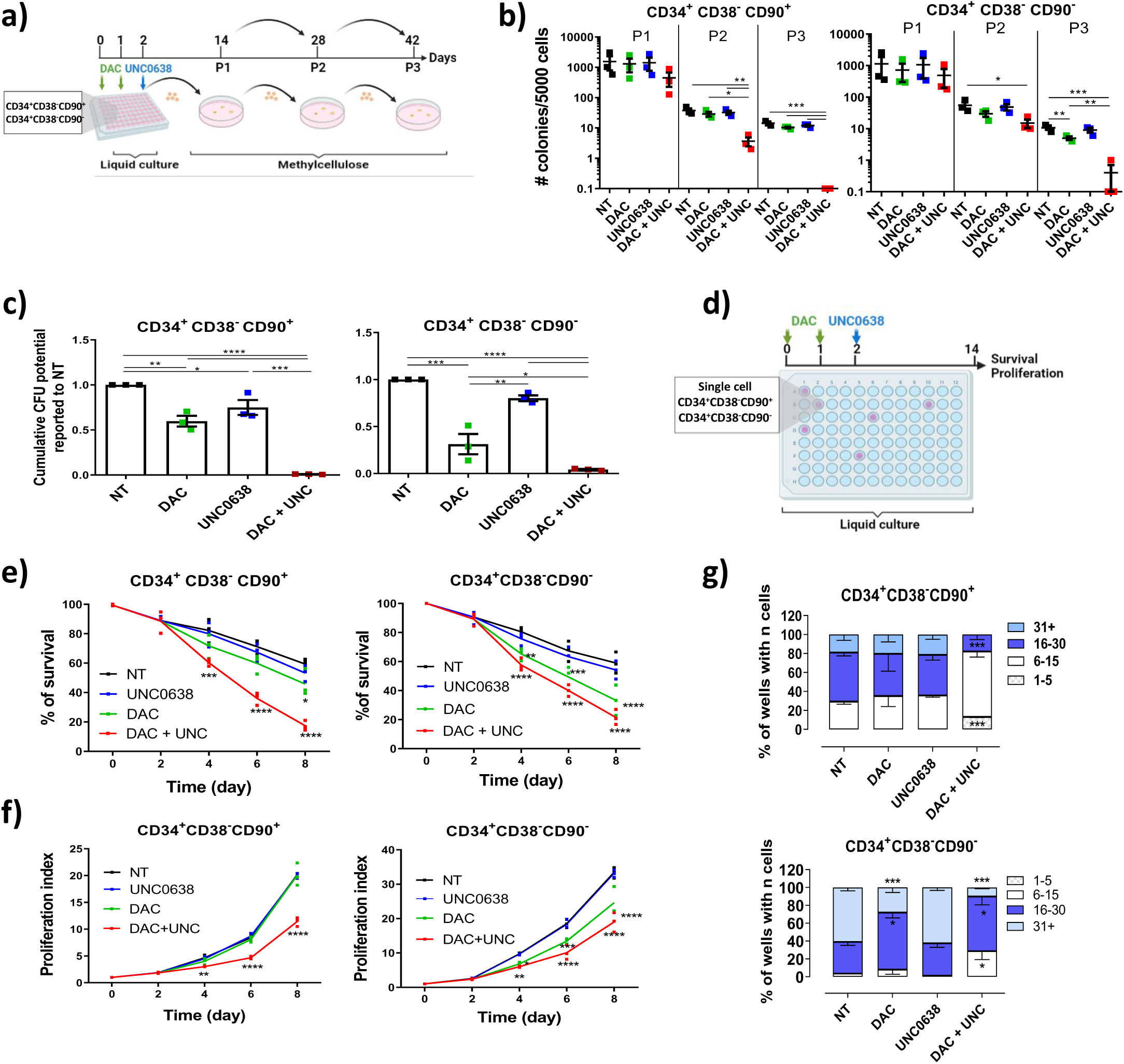
Specific targeting of CMML mutant HSCs by DAC and G9A/GLP inhibitor combination. **a)** Protocol for treatments and serial replating assays in methylcellulose using stem cell- (CD34^+^CD38^-^CD90^+^) or progenitor- (CD34^+^CD38^-^CD90^-^)-enriched populations. P1, passage 1, P2, passage 2, P3, passage 3. **b)** Number of colonies formed from CMML patient stem and progenitor cells (n=3) after the different passages in the presence or absence of DAC +/- UNC0638. Numbers are reported to 5000 cells. Mean +/- SEM. One-way ANOVA. **c)** Effects of DAC and/or UNC0638 treatments on the cumulative clonogenic potential ([number of colonies at P2/number of cells planted at baseline] X number cells retrieved at P1) of stem and progenitor cells from patients, n=3. Mean +/- SEM. One-way ANOVA. **d)** Protocol for treatments and single cell liquid culture assays using stem cell- (CD34^+^CD38^-^ CD90^+^) or progenitor- (CD34^+^CD38^-^CD90^-^)-enriched populations. **e-g)** Single cell liquid cultures. Viability (e), proliferation index (f) and cell number distribution per well at day 8 of the culture (g) of CMML patient stem cells (left panel) or progenitors (right panel) treated with either DAC (green), UNC0638 (blue) or both (red), or left non-treated (NT, black). n=3. Mean +/- SEM. Two-way ANOVA Bonferroni’s Multiple Comparison. *p<0.05; **p<0.01;***p<0.001;****p<0.0001.

We further explored drug effects in single-cell liquid culture experiments for 14 days (Fig. 5d), measuring cell viability (number of wells with at least one cell) and proliferation (total number of cells in wells with at least one cell). In the absence of treatment, viability and expansion of both stem cells and progenitors were similar in control and patient samples (Supplementary Fig. 8a and b). However, the colony size formed by CMML cells was larger than those of controls (Supplementary Fig. 8c). The combination reduced the viability and expansion of both patient stem cells and progenitors (Fig. 5e-g). DAC treatment alone targeted patient progenitors only, UNC0638 alone had no effect, and none of the treatments impaired the viability and proliferation of control stem cells and progenitors (Supplementary Fig. 8d-f).

We subsequently interrogated the ability of the drug combination to selectively eliminate leukemic cells by performing Sanger sequencing of pre-identified mutated genes (Supplementary Table 2) in colonies grown from CD34^+^ cells in methylcellulose and from CD34^+^CD38^-^CD90^+^ cells cultured at one cell per well for 14 days, focusing on somatic mutations with a variant allele frequency (VAF) of 30% or more. Detecting mutations with the highest VAF likely captures the primary events and major clones, as previous studies demonstrated that mutations accumulate almost linearly in CMML stem and progenitor cells, with limited branching events and early clonal dominance^2^. In line with previous findings^2^, a minority of wild-type cells (ranging from 0 to 40%) were found in untreated conditions (Fig. 6). The lack of detection of wild-type cells in patient #3564 could be due to an overrepresentation of mutated clones in methylcellulose^2^ and the limited number of clones analyzed. While DAC and UNC0638 showed limited effects when tested alone, the combination drastically reduced the proportion of mutated clones in all patients (Fig. 6). In patients #3459 and #3562, with mutations in *TET2* and *KRAS*, respectively, the major clones specifically and completely disappeared. Patients #3480 (harboring clones with 0, 1, or 2 mutations) and #3564 (harboring clones with 1, 2, or 3 mutations), showed increased proportions of non-mutated cells, along with reduction in the most mutated clones carrying 2 or 3 mutations. UNC0638 alone slightly increased the proportion of clones or colonies without any mutation (patients #3459 and #3562) or harboring 1 vs. 2 mutations. DAC alone tended to favor the growth of clones harboring mutations (patients #3459 and #3562) or the most mutated clones (increased size of clones with 3 mutations from 59 to 78% in patient #3564). Altogether, these results show that DAC and UNC0638 demonstrate a synergistic and specific impact on leukemic cells while preserving wild-type cells.

**Figure 6:**
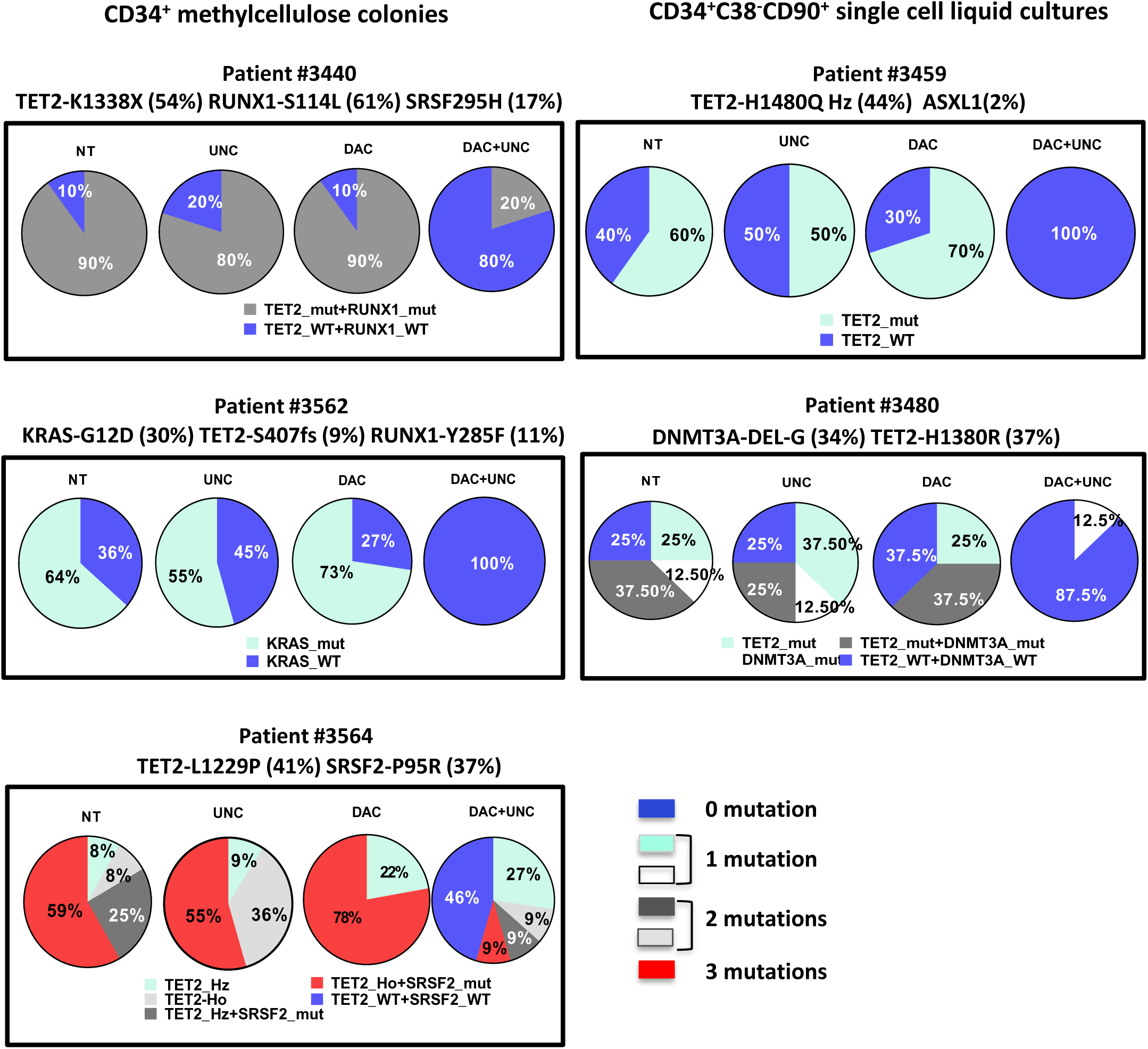
the drug combination selectively eliminates leukemic cells while sparing wild-type cells in the patients. Evaluation of the variant allelic frequency of identified mutated genes with VAF >30%, after 14 days of culture of CD34^+^ cells in methylcellulose (left panel) or CD34^+^CD38^-^CD90^+^ stem cells in liquid cultures (right panel), in the 4 conditions, as indicated. Percentage of cells with 0 (blue), 1 (light green and white), 2 (light and dark grey), or 3 (red) mutations. Mutation VAF are indicated over each panel.

### The HMA and G9A/GLP inhibitor combination increases TE expression in CMML CD34^+^ HSPCs

To assess the impact of DAC and UNC0638 on TE expression, we performed RNA-seq analysis on CMML CD34^+^ cells (n=4 patients) treated in liquid culture with or without DAC and/or UNC0638 (Fig. 7a). Cells were harvested at day 4. At that time, the number of cells was not significantly different between treated and non-treated conditions (Supplementary Fig. 9a). Multimapping analysis of RNA-seq data revealed 340 differentially expressed TE families (p<0.05), all exhibiting increased expression after treatment with the DAC/UNC0638 combination compared to untreated cells (Fig. 7b; Supplementary Data 7). In contrast, only two differential TE families were identified after 4 days of treatment with either DAC or UNC alone (Supplementary Fig. 9b and c). Upon DAC/UNC0638 combination, ERVs and DNA transposons accounted for 43% and 32% of differentially expressed TEs, respectively (Fig. 7c). Notably, ERV retrotransposons showed significant enrichment compared to LINEs and DNA transposons (Fig. 7d). Comparative analysis of these deregulated TEs across all culture conditions revealed that DAC and UNC0638 alone elevated the expression of ERV and DNA transposons compared to untreated cells, with the combination synergistically increasing the expression of ERVs, DNA and LINEs (Fig. 7e; Supplementary 9d). None of the treatments affected SINE expression.

**Figure 7:**
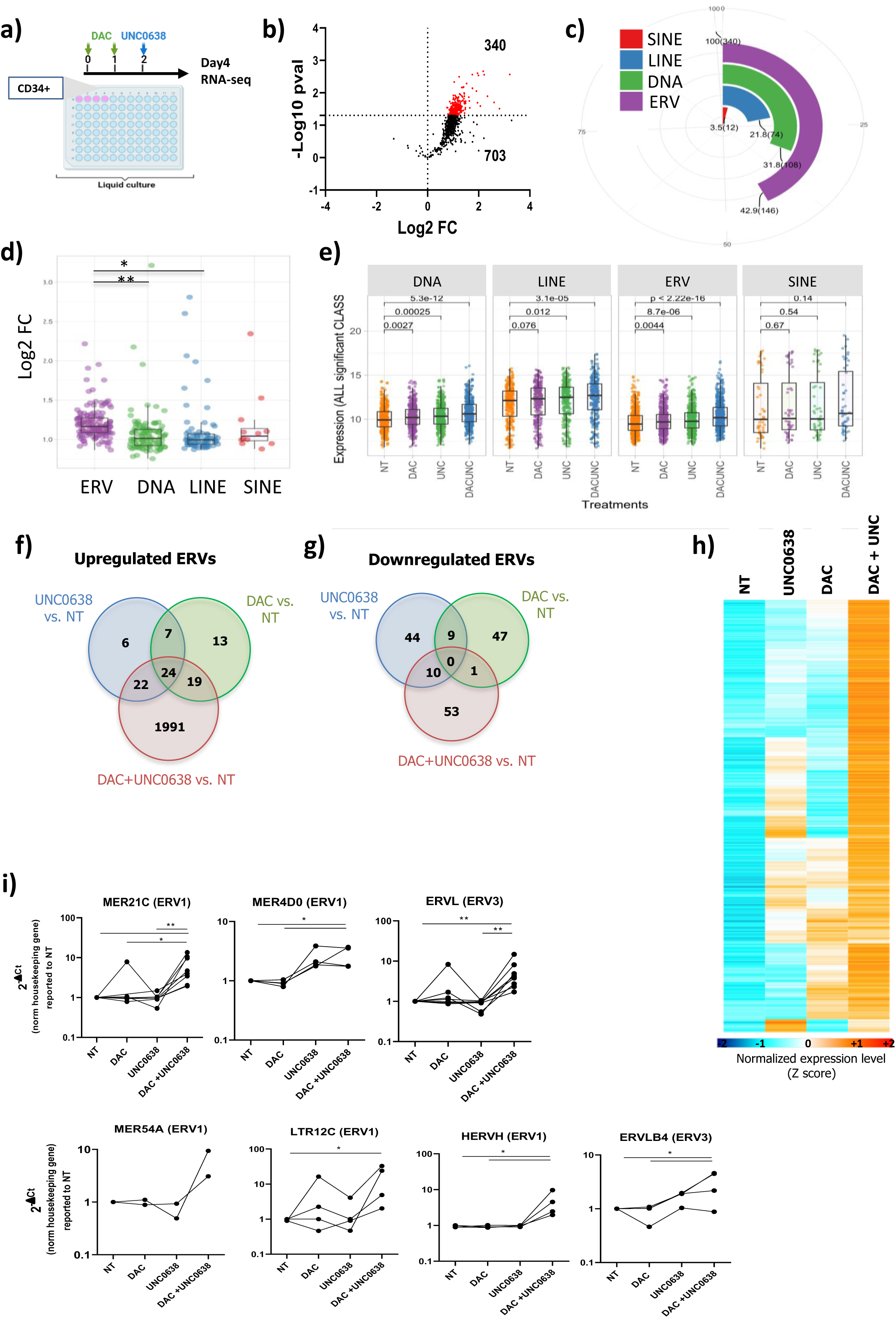
Synergistic increase in TE expression in CMML patients CD34^+^ cells after treatment with the combination. **a-e)** TE expression in RNA-seq analysis from CD34^+^ cells from 4 patients, after treatments in liquid culture with either DAC, UNC0638 or DAC+UNC0368, or left untreated (NT). **a)** Protocol for treatments in liquid culture. **b)** Volcano plots showing differential TE expression (pval<0.05) in the DAC+UNC0638 condition compared to NT cells. **c)** TE family (SINE, LINE, LTR, DNA) repartition in differentially expressed TEs. Numbers indicate the percentage represented by each family. **d)** Log2 Fold-change increase of differentially expressed TE classes in the DAC+UNC0638 condition vs. NT cells. One-way ANOVA Bonferroni’s Multiple Comparison. **e)** Expression levels of TEs differentially expressed in the DAC+UNC0638 vs. NT condition in each of the treatment conditions, sorted by TE class. **f-h)** Differential analysis of ERV loci in RNA-seq data from 7 patients after treatments with DAC, UNC0638 or both compared to the untreated condition (NT). **f, g)** Venn diagrams of the differentially expressed ERV transcripts (pval<0.05, FC 1.5), up- (e) and down-regulated (f) in each of the treated condition vs. NT, as indicated. **h)** Heat map representation of the median expression level of differentially expressed HERVs in DAC+UNC0638 in the different conditions. Clusters with fewer than 20 nodes were excluded from the analysis. **i)** RT-qPCR analysis of mRNA expression of chosen ERV families in the different conditions. Ct were normalized to HPRT, Tubulin, Gus, PPIA and/or RPL32 and reported to the NT condition. One line represents a single patient. Means +/- SEM. One-way ANOVA Tukey’s Multiple Comparison * p<0.05; **p<0.01

Focusing on ERVs that displayed the most significant differential expression upon DAC/UNC0638 treatment, we conducted a loci-based quantification, as described in our previous work^31^, incorporating four additional patient samples. We identified 2,120 differentially expressed ERV loci (p<0.05, FC 1.5) between untreated and DAC/UNC0638 conditions, most of them (97%) upregulated (Fig. 7e and f; Supplementary Data 8). Confirming the results above, the impact of DAC or UNC0638 treatment alone was notably weaker, resulting in 120 and 122 differentially regulated ERV loci, respectively, compared to non-treated condition (Fig. 7f and g). A majority of the ERVs induced by the combination (1991/2056) were specific to this condition, whereas either treatment alone upregulated very few specific ERVs (10% and 20% for UNC0638 and DAC, respectively). Conversely, differentially downregulated loci were largely specific to each condition (48/58 and 43/43 of the ERVs downregulated by UNC0638 or DAC alone). Focusing on ERVs differentially expressed in all conditions compared with untreated cells, the vast majority of loci exhibited a synergistically higher expression level in cells treated with the combination than in DAC or UNC0638 treatment alone (Fig. 7h). RT-qPCR experiments targeting ERV family members deregulated by the combination further validated their greater or synergistic induction after 4 days of treatment with DAC/UNC0648 compared to either inhibitor alone (Fig. 7i). Together, these results indicate that the HMA and G9A/GLP inhibitor combination, but not each inhibitor alone, induces a significant and rapid derepression of TEs, particularly ERVs, in CMML CD34^+^ cells.

### The DAC/UNC0638 combination reactivates viral and inflammatory pathways in CMML cells

ERVs can generate double-stranded RNAs (dsRNAs) that are recognized by antiviral pattern sensors, leading to IFN-I production and activation of the IFN/NFκB pathway—a process known as viral mimicry, which exhibits an anti-tumoral effect^13,16,17^. Analysis of coding gene expression revealed 3,227 DEGs (p<0.05, FC 1.2), of which 2,914 (90%) are positively and 313 (10%) are negatively expressed in DAC/UNC0638 treated compared with untreated CD34^+^ cells (Fig. 8a and b; Supplementary Data 9). Treatment with DAC or UNC0638 alone induced only 187 and 158 DEGs, respectively, with 56% and 53% being overexpressed, and 44% and 47% being downregulated. GSEA revealed that genes upregulated by the DAC/UNC0638 combination were enriched for IFN, NFκB, and other inflammatory pathways, TP53, and apoptosis. In contrast, downregulated genes included those associated with DNA and histone methylation, transcription, and cell proliferation pathways (Fig. 8c and d and Supplementary 10a-d). No specific gene sets were found to be enriched in cells treated with DAC or UNC0638 alone. Gene Set Variation Analysis (GSVA) revealed that the majority of IFN signaling and IFN production signatures were positively enriched as compared with any individual treatment (Fig. 8e). RT-qPCR experiments showed that the combination significantly increased the expression of *IFNB1* mRNA, encoding IFN-β, and TFs such as *IRF9* and *IRF7* involved in IFN signaling (Fig. 8f), confirming RNA-seq results. Altogether, these data demonstrate that TE derepression induced by the combined treatment correlates with the reactivation of viral and inflammatory pathways in CD34^+^ cells from CMML patients.

**Figure 8:**
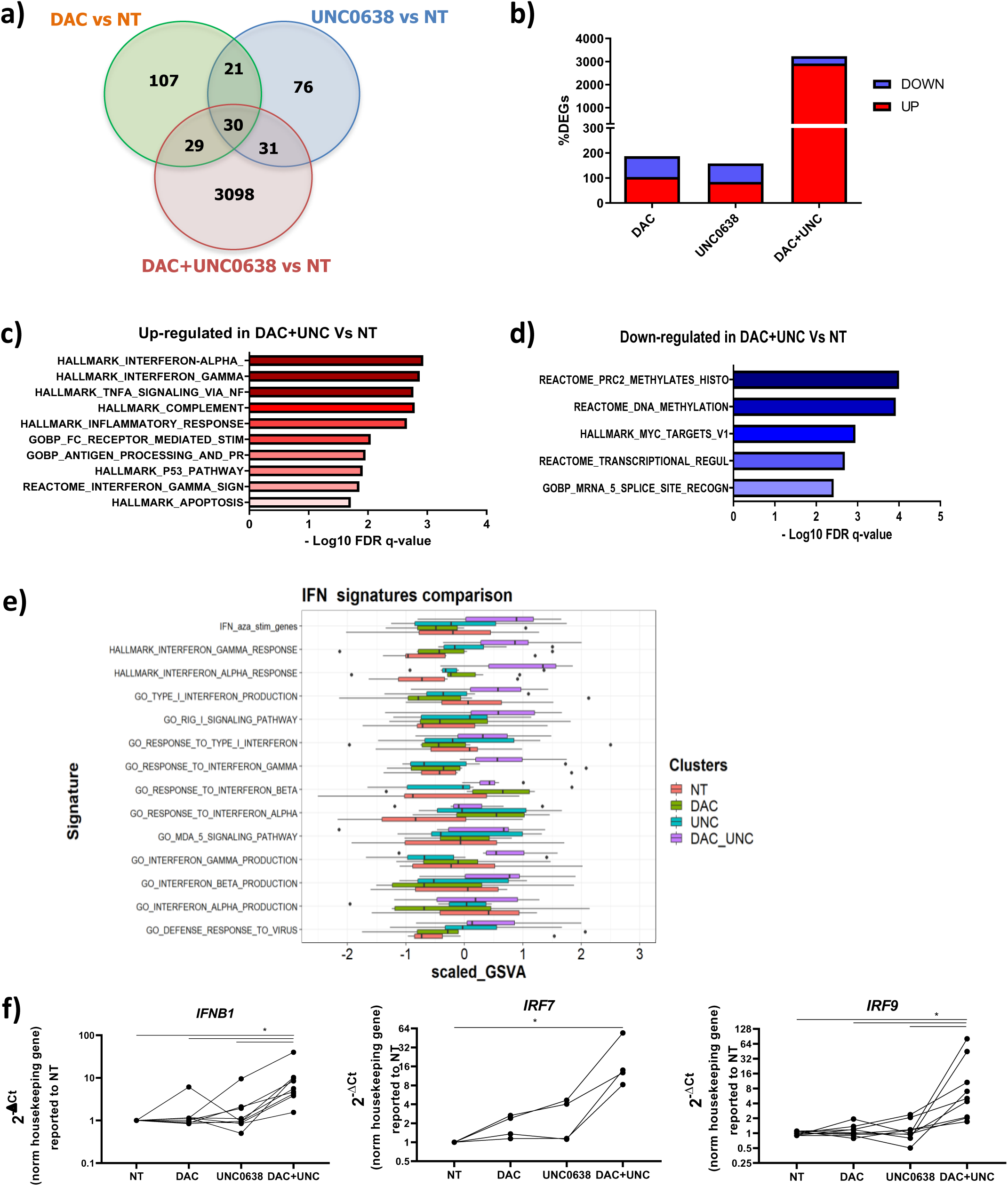
Induction of transcriptional viral mimicry state by the combination of HMA and G9A/GLP inhibitors. **a-e)** RNA-seq-analysis of coding genes from CD34^+^ cells of 7 patients after 4 days of treatment or not with DAC, UNC0638 or the combination. Each condition is compared to NT cells. **a)** Venn diagram showing the number of common or unique DEGs in each condition. **b)** Distribution of DEGs overexpressed (red) and downregulated (blue). **c, d)** GSE analysis of pathways enriched in upregulated (c) and downregulated (d) genes in DAC+UNC0638 vs. NT. **e)** Enrichment analysis of different IFN/antiviral signatures by single sample gene-set variation analysis (ssGSVA). **f)** mRNA expression level of IFNB1, IRF9 and IRF9 measured by RT-qPCR. Ct were normalized by custom housekeeping genes (HPRT, Tubulin, Gus, PPIA and/or RPL32) and reported to the NT condition. Each line represents a single patient. Means +/- SEM. One-way ANOVA Dunnett’s multiple comparison* p<0.05

### The induction of IFN signaling by the combination is essential for eliminating CMML HSPCs

We further examined whether IFN signaling is dysregulated at the protein level. Immunofluorescence analysis demonstrated increased levels of key proteins involved in IFN-I production (IRF3 and IRF7) and IFN signaling (STAT1) in CD34^+^ cells from CMML patients after 4 days of culture with DAC/UNC0638, compared to untreated cells (68-fold, 50-fold, and 3-fold increase, respectively) (Fig. 9a-c). While HMA treatment alone significantly increased IRF7 and STAT1 protein levels, the combination induced a much stronger response. Notably, only the combination led to a substantial increase in the phosphorylated form of STAT1 (pSTAT1) (98-fold compared to untreated), indicating the activation of IFN signaling after 4 days of culture (Fig. 9d). In control cells, only a modest increase in IRF3 was observed upon treatment with the combination (1.5-fold), and this increase was notably lower compared to that observed in patients (68-fold) (Supplementary Fig. 11a-d). These findings further underscore the specific and potent activation of IFN signaling achieved by the combination treatment in CMML CD34^+^ cells.

**Figure 9:**
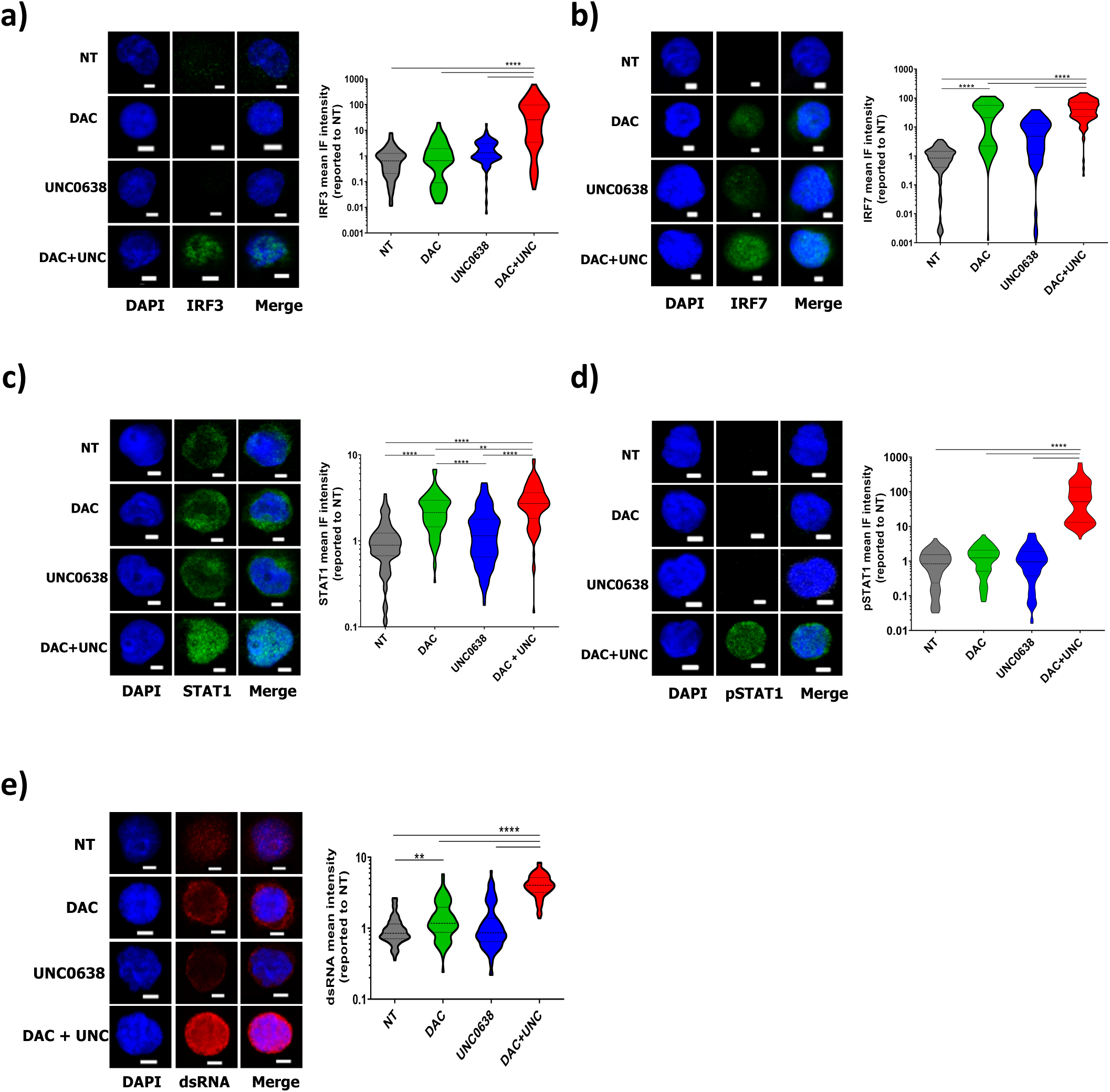
IFN signaling is activated by the HMA and G9A/GLP inhibitor combination. Representative images and quantification of the mean overall immunofluorescence (IF) intensity of **a)** IRF3, **b)** IRF7, **c)** STAT1, **d)** pSTAT1 and **e)** dsRNAs in patient CD34^+^ cells after 4 days of culture. Results are reported to the mean IF intensity in the NT condition. N= 2 patients. Bars, 3 µm. Quantifications were performed using ImageJ software. Violin plots representing the mean IF intensity of each cell. At least 45 cells counted per sample. Dotted lines: median and 25th and 75th percentiles. One-way ANOVA Bonferroni’s Multiple Comparison. **p<0.01;****p<0.0001.

To investigate the potential involvement of ERVs in activating IFN signaling, we assessed dsRNA formation using J2 antibody that specifically recognizes dsRNAs. The combination treatment markedly increased dsRNA expression in patient CD34^+^ cells, while having no significant effect on control cells (Fig. 9e; Supplementary Fig. 11e). Consistent with its modest impact on IRF7 and STAT1 expression, DAC alone showed a slight increase in dsRNA expression, whereas UNC0638 alone had no observable effect (Fig. 9e). We then explored whether cell death induced by the combination could be related to IFN-I signaling activation by treating CD34^+^ cells with the DAC/UNC0638 combination in the presence or absence of an antibody blocking IFNAR2 (IFN-I receptor) before being cultured in methylcellulose (Fig. 10a). Blocking IFNAR2 partially restored the clonogenic capacity of CMML CD34^+^ cells in the presence of DAC/UNC0638 (Fig. 10b and c). This finding highlights the contribution of the antiviral response of malignant cells to the efficacy of the DAC/UNC0638 combination.

**Figure 10:**
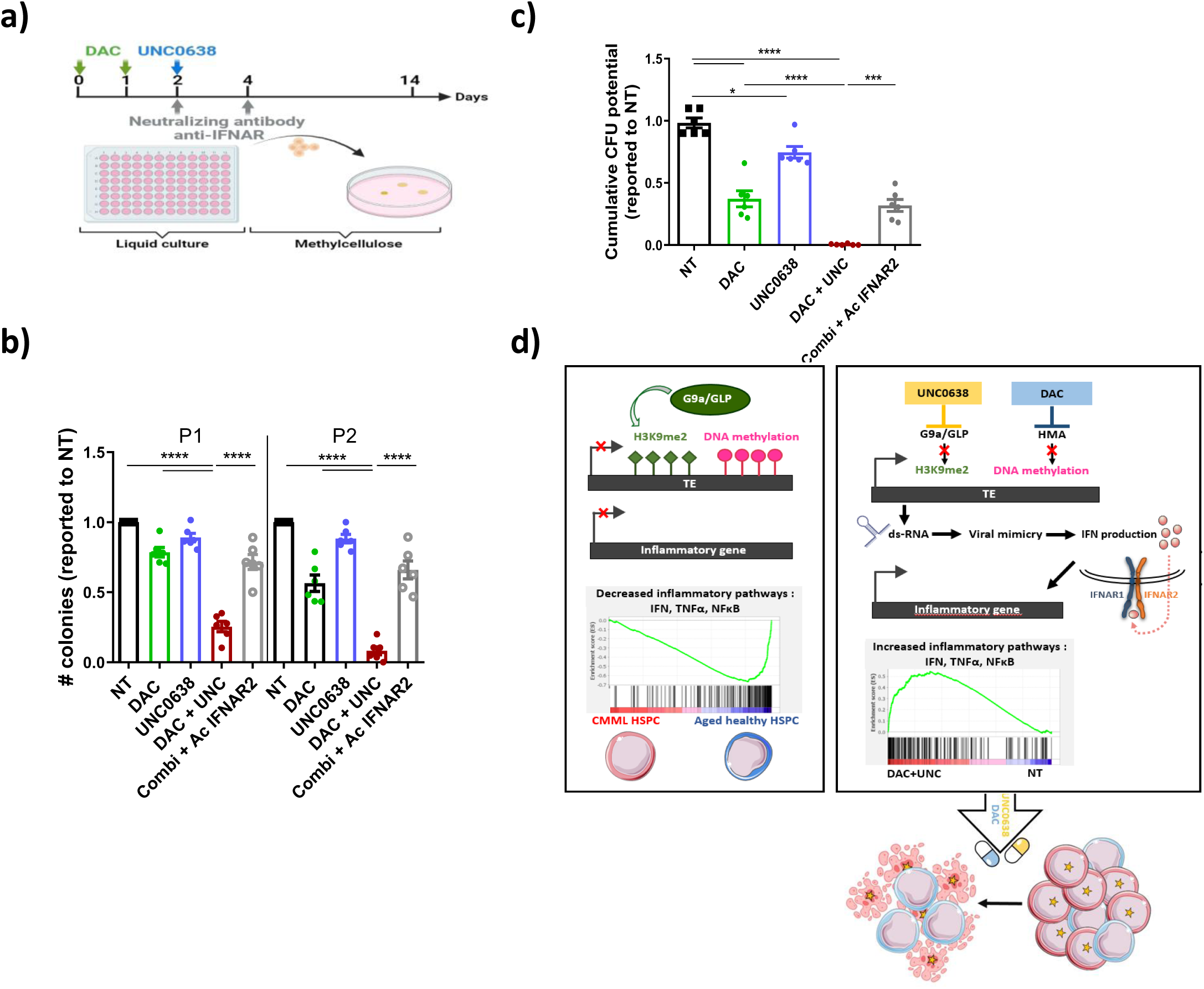
IFN signaling is required for HMA and G9A/GLP inhibitor combination-induced cell death. a) Protocol scheme for the treatments with DAC +/- UNC0638 and blocking IFN receptor (IFNAR2) antibodies. The anti-IFNAR2 antibody (1µg/ml) was added on days 2 and 4 and the cells were seeded in methylcellulose on day 4. b) Number of colonies at passages 1 and 2, reported to the non-treated condition. c) Cumulative potential at passage 2. Results are reported to those obtained in the NT condition. Mean +/- SEM; n=6 patients. One-way ANOVA Bonferroni’s multiple Comparison. p<0.05; **p<0.01;****p<0.0001. d) Model illustrating the increase in H3K9me2 at TEs and repression of immune-associated transcripts in CMML HSPCs compared with age-matched healthy HSPCs (left). The mechanism repressing inflammatory gene is unknown but independent of H3K9me2. TEs are repressed by both H3K9me2 and DNA methylation in CMML cells, as shown by their reexpression only upon treatment with a combination of HMA and G9A/GLP inhibitors. Reexpression of TEs results in dsRNA formation, induction of IFN signaling and reactivation of the innate immune response in CMML HSCs, selectively targeting mutated cells while preserving residual wild-type HSCs. Dark blue cell: healthy donor HSPC. Red cells: CMML HSPC. Red cell with a yellow star: CMML mutated HSC. These cells enter in apoptosis upon treatment with the combination. Light blue cell: non-mutated HSC in CMML patients. Dark blue cell: healthy donor cell.

## Discussion

This study reports the first comprehensive characterization of chromatin and transcriptomic changes in CMML compared to age-matched healthy donor CD34^+^ cells upon *ex vivo* exposure to HMAs. We demonstrate that CMML CD34^+^ cells, regardless of their mutation pattern, exhibit repression of immune and age-associated transcripts, coupled with an increase in the repressive H3K9me2 mark, predominantly at TEs. This chromatin context may protect them from both TE activation^9,14^ and the highly inflammatory environment that characterize aging, and promote their expansion^18,19,22,32,33^. Reactivation of TEs and TE-induced innate immune response, achieved through a combination of HMAs and G9A/GLP inhibitors, appears to decrease the fitness of mutated cells compared to residual wild-type cells (Fig. 10c). This discovery opens up new therapeutic avenues for treating this severe myeloid malignancy.

The depicted downregulation of proinflammatory genes in CMML compared to age-matched healthy donor CD34^+^ cells contrasts with the proinflammatory transcriptome of CMML monocytes^32^ and the increased level of various inflammatory cytokines in peripheral blood plasma of CMML patients^33–35^. HSC aging is associated with an increased expression of proinflammatory genes^18,19,22,36^ likely indicative of the cumulative impact of infections and various stresses over the lifespan, and chronic inflammation promotes premature HSC aging in mice^37^. Recent findings highlight chromatin accessibility changes in aged mouse HSCs, with open regions notably enriched for TF activated in response to cytokine signals and inflammation, such as the JAK-STAT pathway. These open regions exhibit a correlation with gene expression levels^18^. Senescent human HSPCs exhibit heightened expression of TEs and inflammatory genes^38^. The presence of enriched NFκB and IRF motifs in peaks displaying reduced chromatin accessibility suggests that CMML HSPCs may have a diminished capacity to respond to this highly inflammatory milieu. This might protect them from the deleterious effects of aging and inflammation when compared to wild-type CD34^+^ cells, a difference that should contribute to the expansion of the leukemic clone. Supporting this hypothesis, we observed a decline in senescence and aging signatures in CD34^+^ cells from CMML compared to those from aged healthy donors. Our recent results also indicate that dysplastic granulocytes generated by the CMML clone produce high levels of CXCL8 that inhibits the *ex vivo* growth of wild-type CD34^+^ cells while having limited effects on mutated CD34^+^ CMML cells, thereby promoting leukemic clone expansion^39^. Consistent with these findings, studies in zebrafish and humans have reported that the muted inflammatory response of mutant HSCs, compared to wild-type HSCs in the same samples, renders them more resistant to inflammation and aging, facilitating their selection^40,41^. These results suggest that epigenetic drugs that enable the reactivation of inflammatory/antiviral responses in CMML HSCs may break out such vicious circle in CMML as well as other age-associated malignancies.

In a previous work, we showed that HMAs could restore a balanced hematopoiesis in CMML patients without decreasing the mutation VAF nor preventing clonal evolution, indicating that these compounds could not eradicate the malignant clone^4^. Similar observations were made in MDS patients treated with 5-AZA^42,43^. An additional observation is provided by our experiments indicating that HMAs selectively target progenitors while having no detected effect on stem cells in both liquid culture and methylcellulose assays. Even more, DAC appears to slightly promote the growth of the most mutated cells, which could explain why CD34^+^ cells from HMA-responsive MDS patients were observed to be enriched in progenitor cells as compared with non-responders^42,43^. Similarly, HMAs were reported to predominantly target granulo-monocytic progenitors in high-grade MDS and acute myeloid leukemia (AML) ^43^. The high expression of cell cycle-associated genes depicted in CMML CD34^+^ cells who respond to HMAs may promote their incorporation in nucleic acids^44^. The lower expansion of stem cells compared to progenitors (mean fold expansion of 20.34 +/- 0.34 and 32.26 +/- 0.98 in 8 days, respectively) may also contribute to the greater sensitivity of progenitors to HMAs. Alternatively, differences in DNA and histone methylation, expression of genes encoding RNA helicases and autophagy genes^45^ might exist between CMML progenitor and stem cells.

Our findings indicate that DAC as a single agent is an inefficient inducer of either TE expression or inflammatory / IFN signaling in CMML CD34^+^ cells *ex vivo*. The role of these events in HMA activity in myeloid malignancies remains controversial. 5-AZA was initially reported to upregulate the expression of some ERVs in primary CD34^+^ cells collected from MDS patients^46^. Overexpression of TEs in DNMT3A-mutant AML cells renders them more sensitive to 5-AZA via viral mimicry response^47^. An increased transcription of evolutionary young TEs with an activation of innate immune responses was depicted in some 5-AZA-responders^48^ but not validated in others^49^. These discrepancies could be related to the heterogeneity of the studied cohorts with diverse diseases^48,49^ and genetic backgrounds^50,51^ as well as methodological issues.

While TEs are commonly recognized for their tumor-promoting functions, recent studies have suggested that RTEs, including active LINE-1 and ERVs, have been evolutionarily selected for their tumor-suppressing function through their ability to induce viral mimicry and/or p53 activation^52^. Accordingly, TE transcription is commonly suppressed along tumor progression, for example from low-grade to high grade MDS, then a full-blown AML^45^, where a low repeat/gene expression ratio correlates with a poor outcome^53^. This is associated with the inactivation of innate immune pathways, including IFN and NFκB signatures, potentially contributing to the immune evasion mechanism observed in this pathology. One study also reported that CMML and MDS CD34^+^ cells expressed lower levels of TEs than healthy donor cells^49^. Epigenetic mechanisms that were shown to contribute to TE repression in myeloid malignancies include the H3K9 methyltransferase SETDB1^54^ and the epigenetic repressor complex HUSH whose component MMP8 mediates LINE-1 element repression^55^. The decrease in H3K9me3 observed in CMML CD34^+^ cells suggests this mark may not be involved in the repression of TEs and TE-induced innate immune response in these cells. The sharp increase in H3K9me2 and G9A/GLP methyltransferase expression, which was detected in CMML cells may compensate for H3K9me3 loss. A similar increase in H3K9me2 expression was shown to prevent type I IFN pathway activation and immune gene expression in AML cell lines^24^. Our CUT&Tag analysis show that H3K9me2 is mainly located at TEs and did not indicate a direct inhibitory effect of H3K9me2 on immune gene expression. In addition, the G9A/GLP inhibitor UNC0638 alone did not induce IFN/inflammatory signaling in CMML CD34^+^ cells. G9A interacts with DNMT1 and recruits DNMT3A/B through MPP8 to promote the compaction of chromatin in H3K9me2-marked regions^56^. In embryonic stem cells, loss of H3K9me2 was shown insufficient to generate a global DNA demethylation^57^. In HSPCs, the H3K9me2 mark is detected preferentially in demethylated regions and participates in global changes in chromatin structure during myeloid differentiation^27^. This could explain a synergistic effect of G9A/GLP inhibition with HMAs on TE expression.

Besides controlling TE expression, G9A/GLP has been shown to play a role in AML leukemic cell stemness^58^. Altogether, the present study designates G9A/GLP inhibition as a promising strategy to specifically target leukemic stem cells in CMML. Even if the combination slightly impacted control cells *in vitro,* its effect was far more efficient on mutated cells, as shown by the selective eliminatation of mutated leukemic HSCs while preserving wild-type cells in the same patient CMML sample (Fig. 6). Unfortunately, our attempts to explore the ability of HMAs combined to the current G9A/GLP inhibitors to eradicate leukemic stem cells in patient-derived xenografts established in mice was precluded by the toxic side effects of the combination compounds. This could be due to small effect of the combination on healthy murine progenitors. Less-toxic direct inhibitors or alternative indirect approaches will be needed to validate the merit of this therapeutic approach.

## Materials and methods

### Patient and healthy donor samples and processing

The study was carried out on patients diagnosed with CMML, before any treatment according to the 2016 iteration of the WHO classification (Supplementary Table 1). Patient BM samples were obtained as part of a BM assessment of the disease carried out in one of the hematology department affiliated to the Groupe Francophone des Myélodysplasies (GFM). Informed consents were obtained in agreement with Ile-de-France ethics committee (MYELOMONO cohort, DC-2014-2091). Mutations were detected in patients’ monocyte DNA by next generation sequencing (NGS) using a myeloid panel, as previously described ^4^. Age-matched healthy donor BM samples were obtained from femoral heads from individuals who had undergone surgery to treat mechanical osteoarthritis, thanks to the banque de tissus osseux, (Hôpital Cochin). Cord blood was obtained from Unité de Thérapie Cellulaire, CRB-Banque de Sang de Cordon, (AP-HP Hôpital Saint-Louis, Paris) - Authorization no.: AC-2022-5325. Informed consent was obtained from all donors, in accordance with the Declaration of Helsinki. Samples were collected sequentially and cell culture were performed directly on arrival without prior freezing / thawing. Patient samples were collected by BM puncture with EDTA aspiration. BM cells from healthy elderly subjects were obtained by flushing the femoral head in PBS (Gibco). After washing and resuspension in PBS, the cells were filtered on 70 µM nylon mesh. HSPCs were obtained from mononuclear cells sorted on a Ficoll gradient (Pancoll, PAN-Biotech) and isolated by positive magnetic cell sorting performed on CD34 marker expression following the manufacturer’s instructions (AutoMACS ProSeparator, Miltenyi Biotech). For sorting stem cell-enriched (CD34^+^CD38^-^CD90^+^) and progenitor-enriched (CD34^+^CD38^-^CD90^-^) populations, cells were further stained with CD38-PE-Cy7 (BD Biosciences; clone HB-7), CD90-FITC (Sony Biotechnologies; clone HIT2) and sorted on ARIA fusion cell sorter.

### Cell cultures and treatments

Freshly isolated human HSPCs were plated at 200,000 cells/ml and cultured in IMDM medium supplemented with 25% BIT9500 (StemCell Technologies), 1% penicillin/streptomycin (P/S), containing the following cytokines (all from Miltenyi Biotech): FTL3-Ligand (FTL3-L, 100ng/μL), Thrombopoietin (THPO, 100ng/μL), stem cell factor (SCF, 60ng/μL) and interleukin 3 (IL3, 10ng/μL), at 37°C, 20% O2, 5% CO2. For treatments, decitabine (DAC, Sigma Aldrich), or 5-Azacytidine (AZA, Sigma Aldrich) were added at the start of the cultures (day 0) and again on day 1. G9A/GLP inhibitor (UNC0638 or UNC0642, MedChem) was added once on day 2. The dual DNMT/G9A inhibitor, CM-272 (Axon Medchem) was added on day 0 and again 48h later and the cells were directly seeded in methylcellulose.

### Single cell colony and CFU assays

Single human or mouse HSCs/HSPCs were FACS-sorted into individual wells of round-bottom 96 well plates and cultured in the presence or absence of inhibitors. At different times starting day one, each well was checked manually for the the presence of cells. For CFU assays, 300 CD34^+^ or 100 CD34^+^CD38^-^CD90^-/+^ treated or not with the inhibitors were seeded in duplicate in 24-well plates containing 300µl of methylcellulose (H4230, StemCell), supplemented with 1% P/S (ThermoFischer Scientific), FLT3-L (25ng/µL), THPO (25ng/µL), SCF (60ng/µL), IL3 (10ng/mL), EPO (2 units), GM-CSF (10ng/mL). Colonies were counted after 14 days of culture. The mean size of the colonies was estimated by harvesting them and counting the total number of cells in the well. For replating assays, 5000 cells were replated in methylcellulose in the same medium. This process was repeated twice. The cumulative CFU potential that takes into account both the number of CFUs and their size was calculated at passage 2 described ^30^.

### Gene mutation screening in colonies

Mutations with VAF > 30% were analyzed after *in vitro* treatment (Supplementary Table 2).

Colonies from CD34^+^ or CD34^+^CD38^-^CD90^+^ cells grown in methylcellulose or in single cell liquid cultures, respectively, in the presence or absence of DAC and/or UNC0638, were picked up individually after 14 days, washed and stored at -80°C as dry pellets. Cells were lysed with a solution containing proteinase K (0.1 mg/mL, Fisher Scientific), 2% Tween-20 (Sigma-Aldrich) and incubated for 1 h at 65°C followed by 15 min at 95°C. DNA was amplified by PCR using the Q5 High-Fidelity DNA polymerase kit (New England BioLabs) and primers designed around the mutations (Supplementary Table 3). Amplicon sizes were verified by migrating the amplified DNA sample onto a 1.8% agarose gel. The amplified DNA fragments were analyzed by Sanger sequencing (Eurofins Genomics, Ebersberg, Germany). 8-12 clones were tested per condition.

### ELISA for DNA methylation

Cell lysis and extraction of genomic DNA from dry pellets were performed using the NucleoSpin Tissue XS kit (Macherey-Nagel). Genomic DNA quality and quantity were measured using the NanoDrop (ND-1000, Marshall Scientific). Global DNA methylation was studied using the global DNA methylation assay kit - LINE-1 (Active Motif) according to the manufacturer’s instructions.

### Immunofluorescence

5000 human and murine HSPCs/HSCs were fixed with 4% paraformaldehyde (PFA) for 10 min at RT and centrifuged (Tharmac) on poly-L-Lysine slides (Fisher Scientific). Immunofluorescence was performed as described previously^11,14^. The primary antibodies used are listed in Supplementary Table 4. Detection was performed using anti-rabbit secondary antibody coupled to Alexa-Fluor 488 or anti-mouse coupled to Alexa-Fluor-555. All slides were viewed under a TCS SPE confocal microscope (Leica). Images were analyzed and fluorescence intensities calculated using ImageJ software, as described^11,14^.

### RNA purification and quantitative RT-PCR

50,000 - 150,000 cells were lysed in Tri-Reagent solution (Zymo Research) and stored at -80°C until use. Total RNA was extracted using the Direct-Zol RNA microprep kit (Zymo Research). RNAs were reverse transcribed using the EZ DNAse VILO IV kit (Invitrogen). 1.25 µL of cDNA was pre-amplified for 15-18 cycles in a multiplex reaction using Preamp Master-Mix solution (ref. 100-5580, Fluidigm) and primer mix (200 µM of each primer; supplemental Table S3), as described^11^. Quantitative PCR (qPCR) was performed using LUNA Universal qPCR Master Mix (New England Biolabs) on the 7500 real-time PCR machine (Applied Biosystems). Data were normalized with the mean expression of household genes (HPRT, Tubulin, GUS, PPIA and / or RPL32).

### ATAC-seq and data processing

100,000 freshly isolated CD34^+^ cells were processed with the ATAC-seq kit (Active Motif). Transposed DNA fragments were amplified 10-fold by PCR using adapters supplied by the kit. PCR purification was carried out using MinElute PCR Purification Kit (Qiagen, 28004) to remove remaining primers and large fragments. The quantity and quality of the libraries were assessed on Agilent 2100 Bioanalyzer (Agilent Technologies 50567-4626). Sequencing of the libraries was performed on the NovaSeq-6000 at Gustave Roussy (Illumina; 50 bp paired-end reads). ATAC-seq data were processed essentially as described^15^. Read quality was assessed using Fastqc^59^ (v0.11.9) and MultiQC^60^. Bowtie2^61^ (v.2.3.4.1) using the «--very-sensitive» parameter was used for aligning ATAC-seq reads to the human genome version GRCh38/hg38 and Samtools (v.1.7) for data filtering and file format conversion. Duplicate reads were removed with Picard tool - MarkDuplicates (http://broadinstitute.github.io/picard), blacklist regions, chrM and Y were removed with BEDtools intersect and grep -v respectively before peak calling. All filtered.bam files were converted to bedgraphs using the DeepTools bamCoverage subcomm and, with the reads per kilobase of transcript, per million mapped reads (RPKM) normalization method. MACS2 (v.2.1.1)^62^ algorithm was used for peak identification (Q-value cut-off = 0.01). Gained and lost peaks were identified from the narrow peaks in two steps. First, the peak lists from control and CMML patients were merged separately with the subcommands cat and mergeBed to obtain consensus peaks. Second, with intersectBed and the parameters --a <consensus peaks> --b <narrow peaks> --wa --u and subsequently --v, we compare and subtract them, to take the gained and lost peaks for each group. The reads of these consensus peaks were counted and a statistical model based on edgeR^63^ was used to identify the significantly differential peaks. To identify the common peaks between patients or controls, DESeq which uses the Negative Binomial distribution to compute a P value and a fold change for each estimated peak was used. Peaks highly enriched in comparison to the rest were considered gained. Annotation of peaks to genes (100 kb upstream and 25 kb downstream from the TSS) and genomic distribution of accessible regions identified by MACS2 was performed using BEDTools and the -closestBed and -intersectBed subcommands, respectively. Clustering of regions was generated with the ComputeMatrix function of DeepTools, using the reference point --referencePoint center -b 2500 -a 2500 -R <bed files> - s <bigwig files> as parameters. The function plotHeatmap from the same package was used for displaying the average profiles heat map. Differentially accessible chromatin regions were scanned for enriched short-sequence motifs using HOMER software ^64^ with “findMotifsGenome.pl” command.

### RNA-sequencing and data processing

RNA quality was checked on the Agilent Bioanalyzer and quantified. 100-250 pg of total RNA extracted from CD34^+^ cells were used to prepare cDNA libraries, as described^11^, using the SMARTer Ultra Low Input RNA Kit for Sequencing - v3 or -SMARTer Stranded Total RNA-seq kit v3, according to the manufacturer’s recommendations (Takara). Libraries were sequenced in (2 x 100bp) on the NovaSeq-6000 (Illumina). Read quality was assessed with Fastqc v0.11.9, and corrected with Fastqp^59^ v0.23.1. Alignments were performed with STAR (v2.7.10b) on GRCh38 human genome reference. Mapped reads underwent filtration to retain only those associated with regular chromosomes (1-23, XY) and the remaining reads were sorted and indexed. Raw count table were generated with FeatureCounts (package subread, v2.0.1). Transcript expression levels were aggregated in gene expression levels using tximport Bioconductor package v1.13.16. For the CMML versus controls, a batch effect correction was first done using ComBat-seq function of the sva package, before subsequent DESeq2 analysis^65^. PCA analysis was performed with DESeq2, after regularized log transformation of gene counts using default parameters. Pathway analysis were performed and plotted on R using msigdbr (7.5.1), and clusterProfiler (4.2.2 or 4.6) or the GSEA app of MSigDB. GSEA analyses of CMML vs. controls cells were performed and genes pre-ranked on the statistical result of the differential analysis, and pathways were sorted on their normalized enrichment score (NES) before plotting.

Multimapping and expression of TE families was analyzed with TEtranscript (v 2.2.1)^66^. Paired-end reads were aligned to GRCh38 genome using STAR (v2.7.11a) with these parameters: --winAnchorMultimapNmax 100 --outFilterMismatchNmax 3 --alignEndsType EndToEnd -- alignIntronMax 1 --alignMatesGapMax 350. Differentially expressed TEs were identified using the R (v4.1.2) package DESeq2(v1.3.4), with criteria including p-value < 0.05 and an absolute value of log2(FC) > 0. ERV expression was quantified using a loci-based approach, as previously described ^31^. Briefly, RNA-seq reads were aligned to a custom transcriptome using bowtie2 v2.2.1 with custom parameters to retain multimapped reads (-k 100 --very-sensitive-local -- score-min “L,0,1.6”). The custom transcriptome consisted in the hg38 reference transcriptome with 14,968 ERV transcriptional units compiled from RepeatMasker annotations ^67^. ERV and gene expression were then quantified using Telescope^67^ and HTSeq 0.12.3^68^, respectively. Raw counts were concatenated and normalized independently for each dataset using DESeq2 v1.28.0 with variance stabilizing transformation. Differential expression analysis was performed using DESeq2. Raw counts of ERVs and genes were merged and integrated into the same DESeq object, using patient ID and condition as a covariate in the design formula to account for data pairing (design = ∼ patient_id + condition). For each comparison, the untreated (NT) condition was considered as reference. Fold change were shrunk with the ASHR method ^69^. Union of differentially expressed coding genes or ERV were selected based on a 1.5-fold change and pvalue < 0.05. IFN signatures were retrieved from The Molecular Signatures Database (MSigDb). Individual enrichment scores were calculated from each patient by single sample gene-set variation analysis (ssGSVA), and center-scaled.

### CUT&Tag and data processing

CUT&Tag assays were performed using CUT&Tag-IT assay kit (Active Motif) on 50,000 CD34^+^ cells, as described ^11^, using 0.5µg of H3K9me2 antibody (Abcam Ab1220). Sequencing of the libraries was performed on the NovaSeq-6000 (Illumina; 2x 50 bp). CUT&Tag data were processed and analyzed mostly as described^11^. Paired-end reads were aligned to GRCh37 genome using Bowtie2 (v2.4.1) with the following parameters: --end-to-end --very-sensitive --no-mixed --no-discordant --phred33 -I 10 -X 700. Duplicate reads were excluded using Picard (v2.26.9). Reads aligned on regular chromosomes were retained and the aligned read quality score was set to «0» using samtools (v1.13) to get both uniquely mapped and multimapped reads. The Y chromosome was removed from the downstream analysis. The aligned reads underwent two processing methods: 1) direct sorting and indexing BAM files using samtools (v1.13), followed by peak calling with MACS2 (v2.2.7.1) with the specified parameters adapted to broad marks: -B --broad --broad-cutoff 0.1 -f BAMPE -g hs --max-gap 2000 --min-length 200. 2) Downsampling BAM files to the minimum read count across BAM files, subsequent sorting, and indexing using samtools (v1.13). Processed BAM files were then utilized for extracting read counts within peaks. Subsequent downstream analysis was carried out using custom scripts in R (v4.1.2). A matrix of counts for union peaks was generated using chromVAR (v1.16.0) R package and differentially expressed peaks between CMML Patients and controls were identified using the R (v4.1.2) package DESeq2(v1.3.4), with criteria including an p-value < 0.05 and an absolute value of log2(FC) > 0. Peaks consistently identified across all control or patient biological replicates were kept for further analysis. Enrichment of TEs in peaks was performed using a permutation test conducted independently for each TE class. A peak was considered associated with a TE if there is at least 1 base pair overlap between them. Peaks that intersected with any TE belonging to a given TE class were counted in CMML unique (gained) and control unique (lost) peaks. An equal number of peaks were randomly chosen from peaks that do not belong to any section of the Venn diagram, and the count of peaks overlapping with the designated TE class was determined. This iterative process was repeated 1000 times to establish a null distribution for the number of peaks overlapping with the TEs.

Enrichment or depletion of the different classes of TEs was calculated as the log2 fold change of the number of specific peaks overlapping with TEs in any TE class found in patient gained or lost peaks over the median number of TEs of the same class in the randomly selected peaks. The empirical p-value was calculated by dividing by 1000 the number of occurrences where the permutation resulted in values exceeding the number of gained peaks overlapping the TE class.

### Statistics and reproducibility

Results were statistically evaluated by one-way or two-way ANOVA tests or paired and unpaired t-tests using Prism GraphPad PrismTM software (GraphPad Software Inc., San Diego, CA, USA). Graphs show mean and standard error to the mean (SEM). Values of *p<0.05* are considered statistically significant. The use of statistical test, the *p* value, the numbers of samples and experiments are indicated in the respective Fig ure legends.

## Data availability

All raw sequencing Fasq data for RNA-seq, CUT&tag and ATAC-seq presented in this paper will be available through the European Genome-phenome Archive (EGA) under the accession number EGAS50000000367. The data supporting the finduinds of these study are available in Supplementary Data files 1 to 8 joined with the paper. All source data underlying the graphs and charts presented in the main figures are provided in the Supplementary Data 9.

## Supplemental material

Supplementary material includes 11 figures related to Figures 1 to 10, 4 tables and 10 Data files.

## Acknowledgments

We thank Thorsten Braun (Clinical Hematology Department, Hôpital Avicenne, Bobigny), Céline Berthon (Clinical Hematology Department, University Hospital, Lille), Gabriel Etienne (Clinical Hematology Department, University Hospital, Bordeaux), Pierre Fenaux (Hematology Department, Saint-Louis Hospital, Paris) Christophe Willekens (Clinical Hematology Department, Gustave Roussy Cancer Center, Villejuif) for providing patient samples; animal facility, the Genomic and the Imaging and Cytometry Platforms of Gustave Roussy for sequencing and cell sorting and confocal analysis, respectively and Dr. Louis Paré (Nantes University, École Centrale Nantes) for RNA-seq analysis. This work was supported by INSERM and grants from Ligue Nationale Contre le Cancer (LNCC, Equipe labellisée EL2020) and Institut National du Cancer (PLBIO N°2020-095) to F.P. D.H. and A.P. are recipients of fellowship from the Ministère de l’Enseignement Supérieur de la Recherche et de l’Innovation.

## Author contributions

DH, AP MB, AI performed experiments, acquired and analyzed the results. RC, AiP, VA, SD, LL performed bioinformatic analyses. ND performed next generation sequencing and analyzed data. AR, DSB, ET analyzed data. SP, RI provided samples. MM processed samples. ES analyzed data and wrote the manuscript. FPu designed and supervised the study, conducted experiments, analyzed the results and wrote the manuscript.

## Notes

The authors declare no potential conflicts of interest

### Competing Interest Statement

The authors have declared no competing interest.

### Summary of Updates

This version of the manuscriot has been revised by adding new data according reveiwer's request.

## References

1 Khoury, J. D. et al. The 5th edition of the World Health Organization Classification of Haematolymphoid Tumours: Myeloid and Histiocytic/Dendritic Neoplasms. Leukemia 36, 1703–1719 (2022). 10.1038/s41375-022-01613-1

2 Itzykson, R. et al. Clonal architecture of chronic myelomonocytic leukemias. Blood 121, 2186–2198 (2013). 10.1182/blood-2012-06-440347

3 Binder, M. et al. Oncogenic gene expression and epigenetic remodeling of cis-regulatory elements in ASXL1-mutant chronic myelomonocytic leukemia. Nature communications 13, 1434 (2022). 10.1038/s41467-022-29142-6

4 Merlevede, J. et al. Mutation allele burden remains unchanged in chronic myelomonocytic leukaemia responding to hypomethylating agents. Nature communications 7, 10767 (2016). 10.1038/ncomms10767

5 Palomo, L. et al. DNA methylation profile in chronic myelomonocytic leukemia associates with distinct clinical, biological and genetic features. Epigenetics 13, 8–18 (2018). 10.1080/15592294.2017.1405199

6 Solary, E. & Itzykson, R. How I treat chronic myelomonocytic leukemia. Blood 130, 126–136 (2017). 10.1182/blood-2017-04-736421

7 Tsurumi, A. & Li, W. X. Global heterochromatin loss: a unifying theory of aging? Epigenetics 7, 680–688 (2012). 10.4161/epi.20540

8 Zhang, W., Qu, J., Liu, G. H. & Belmonte, J. C. I. The ageing epigenome and its rejuvenation. Nature reviews. Molecular cell biology 21, 137–150 (2020). 10.1038/s41580-019-0204-5

9 Djeghloul, D. et al. Age-Associated Decrease of the Histone Methyltransferase SUV39H1 in HSC Perturbs Heterochromatin and B Lymphoid Differentiation. Stem Cell Reports 6, 970–984 (2016). 10.1016/j.stemcr.2016.05.007

10 Keenan, C. R. et al. Extreme disruption of heterochromatin is required for accelerated hematopoietic aging. Blood 135, 2049–2058 (2020). 10.1182/blood.2019002990

11 Pelinski, Y. et al. NF-kappaB signaling controls H3K9me3 levels at intronic LINE-1 and hematopoietic stem cell genes in cis. The Journal of experimental medicine 219 (2022). 10.1084/jem.20211356

12 Chuong, E. B., Elde, N. C. & Feschotte, C. Regulatory activities of transposable elements: from conflicts to benefits. Nature reviews. Genetics 18, 71–86 (2017). 10.1038/nrg.2016.139

13 Gazquez-Gutierrez, A., Witteveldt, J., S, R. H. & Macias, S. Sensing of transposable elements by the antiviral innate immune system. RNA 27, 735–752 (2021). 10.1261/rna.078721.121

14 Barbieri, D. et al. Thrombopoietin protects hematopoietic stem cells from retrotransposon-mediated damage by promoting an antiviral response. The Journal of experimental medicine 215, 1463–1480 (2018). 10.1084/jem.20170997

15 Clapes, T. et al. Chemotherapy-induced transposable elements activate MDA5 to enhance haematopoietic regeneration. Nature cell biology 23, 704–717 (2021). 10.1038/s41556-021-00707-9

16 Chiappinelli, K. B. et al. Inhibiting DNA Methylation Causes an Interferon Response in Cancer via dsRNA Including Endogenous Retroviruses. Cell 162, 974–986 (2015). 10.1016/j.cell.2015.07.011

17 Roulois, D. et al. DNA-Demethylating Agents Target Colorectal Cancer Cells by Inducing Viral Mimicry by Endogenous Transcripts. Cell 162, 961–973 (2015). 10.1016/j.cell.2015.07.056

18 Itokawa, N. et al. Epigenetic traits inscribed in chromatin accessibility in aged hematopoietic stem cells. Nature communications 13, 2691 (2022). 10.1038/s41467-022-30440-2

19 Chen, Z. et al. Cohesin-mediated NF-kappaB signaling limits hematopoietic stem cell self-renewal in aging and inflammation. The Journal of experimental medicine 216, 152–175 (2019). 10.1084/jem.20181505

20 Grigoryan, A. et al. LaminA/C regulates epigenetic and chromatin architecture changes upon aging of hematopoietic stem cells. Genome biology 19, 189 (2018). 10.1186/s13059-018-1557-3

21 Kamminga, L. M. et al. The Polycomb group gene Ezh2 prevents hematopoietic stem cell exhaustion. Blood 107, 2170–2179 (2006). 10.1182/blood-2005-09-3585

22 Adelman, E. R. et al. Aging Human Hematopoietic Stem Cells Manifest Profound Epigenetic Reprogramming of Enhancers That May Predispose to Leukemia. Cancer discovery 9, 1080–1101 (2019). 10.1158/2159-8290.CD-18-1474

23 Fang, T. C. et al. Histone H3 lysine 9 di-methylation as an epigenetic signature of the interferon response. The Journal of experimental medicine 209, 661–669 (2012). 10.1084/jem.20112343

24 Hansen, A. M. et al. H3K9 dimethylation safeguards cancer cells against activation of the interferon pathway. Science advances 8, eabf8627 (2022). 10.1126/sciadv.abf8627

25 Avgustinova, A. et al. Repression of endogenous retroviruses prevents antiviral immune response and is required for mammary gland development. Cell stem cell 28, 1790–1804 e1798 (2021). 10.1016/j.stem.2021.04.030

26 Liu, M. et al. Dual Inhibition of DNA and Histone Methyltransferases Increases Viral Mimicry in Ovarian Cancer Cells. Cancer research 78, 5754–5766 (2018). 10.1158/0008-5472.CAN-17-3953

27 Schones, D. E., Chen, X., Trac, C., Setten, R. & Paddison, P. J. G9a/GLP-dependent H3K9me2 patterning alters chromatin structure at CpG islands in hematopoietic progenitors. Epigenetics & chromatin 7, 23 (2014). 10.1186/1756-8935-7-23

28 Chou, T. C. Theoretical basis, experimental design, and computerized simulation of synergism and antagonism in drug combination studies. Pharmacological reviews 58, 621–681 (2006). 10.1124/pr.58.3.10

29 San Jose-Eneriz, E., et al. Discovery of first-in-class reversible dual small molecule inhibitors against G9a and DNMTs in hematological malignancies. Nature communications 8, 15424 (2017). 10.1038/ncomms15424

30 Higa, K. C. et al. Chronic interleukin-1 exposure triggers selection for Cebpa-knockout multipotent hematopoietic progenitors. The Journal of experimental medicine 218 (2021). 10.1084/jem.20200560

31 Alcazer, V. et al. HERVs characterize normal and leukemia stem cells and represent a source of shared epitopes for cancer immunotherapy. American journal of hematology 97, 1200–1214 (2022). 10.1002/ajh.26647

32 Franzini, A. et al. The transcriptome of CMML monocytes is highly inflammatory and reflects leukemia-specific and age-related alterations. Blood advances 3, 2949–2961 (2019). 10.1182/bloodadvances.2019000585

33 Niyongere, S. et al. Heterogeneous expression of cytokines accounts for clinical diversity and refines prognostication in CMML. Leukemia 33, 205–216 (2019). 10.1038/s41375-018-0203-0

34 Pronier, E. et al. Macrophage migration inhibitory factor is overproduced through EGR1 in TET2(low) resting monocytes. Communications biology 5, 110 (2022). 10.1038/s42003-022-03057-w

35 Sevin, M. et al. Cytokine-like protein 1-induced survival of monocytes suggests a combined strategy targeting MCL1 and MAPK in CMML. Blood 137, 3390–3402 (2021). 10.1182/blood.2020008729

36 Mann, M. et al. Heterogeneous Responses of Hematopoietic Stem Cells to Inflammatory Stimuli Are Altered with Age. Cell reports 25, 2992–3005 e2995 (2018). 10.1016/j.celrep.2018.11.056

37 Esplin, B. L. et al. Chronic exposure to a TLR ligand injures hematopoietic stem cells. Journal of immunology 186, 5367–5375 (2011). 10.4049/jimmunol.1003438

38 Capone, S. et al. Senescent human hematopoietic progenitors show elevated expression of transposable elements and inflammatory genes. Experimental hematology 62, 33–38 e36 (2018). 10.1016/j.exphem.2018.03.003

39 Deschamps, P. et al. CXCL8 secreted by immature granulocytes inhibits wildtype hematopoiesis in chronic myelomonocytic leukemia. bioRxiv, 2024.2003.2008.583935 (2024). 10.1101/2024.03.08.583935

40 Avagyan, S. et al. Resistance to inflammation underlies enhanced fitness in clonal hematopoiesis. Science 374, 768–772 (2021). 10.1126/science.aba9304

41 Jakobsen, N. A. et al. Selective advantage of mutant stem cells in clonal hematopoiesis occurs by attenuating the deleterious effects of inflammation and aging. bioRxiv, 2023.2009.2012.557322 (2023). 10.1101/2023.09.12.557322

42 Unnikrishnan, A. et al. Integrative Genomics Identifies the Molecular Basis of Resistance to Azacitidine Therapy in Myelodysplastic Syndromes. Cell reports 20, 572–585 (2017). 10.1016/j.celrep.2017.06.067

43 Ali, A. et al. Granulomonocytic progenitors are key target cells of azacytidine in higher risk myelodysplastic syndromes and acute myeloid leukemia. Leukemia 32, 1856–1860 (2018). 10.1038/s41375-018-0076-2

44 Meldi, K. et al. Specific molecular signatures predict decitabine response in chronic myelomonocytic leukemia. The Journal of clinical investigation 125, 1857–1872 (2015). 10.1172/JCI78752

45 Colombo, A. R. et al. Suppression of Transposable Elements in Leukemic Stem Cells. Scientific reports 7, 7029 (2017). 10.1038/s41598-017-07356-9

46 Tobiasson, M. et al. Comprehensive mapping of the effects of azacitidine on DNA methylation, repressive/permissive histone marks and gene expression in primary cells from patients with MDS and MDS-related disease. Oncotarget 8, 28812–28825 (2017). 10.18632/oncotarget.15807

47 Scheller, M. et al. Hotspot DNMT3A mutations in clonal hematopoiesis and acute myeloid leukemia sensitize cells to azacytidine via viral mimicry response. Nat Cancer 2, 527–544 (2021). 10.1038/s43018-021-00213-9

48 Ohtani, H. et al. Activation of a Subset of Evolutionarily Young Transposable Elements and Innate Immunity Are Linked to Clinical Responses to 5-Azacytidine. Cancer research 80, 2441–2450 (2020). 10.1158/0008-5472.CAN-19-1696

49 Kazachenka, A. et al. Epigenetic therapy of myelodysplastic syndromes connects to cellular differentiation independently of endogenous retroelement derepression. Genome medicine 11, 86 (2019). 10.1186/s13073-019-0707-x

50 Bejar, R. et al. TET2 mutations predict response to hypomethylating agents in myelodysplastic syndrome patients. Blood 124, 2705–2712 (2014). 10.1182/blood-2014-06-582809

51 Duchmann, M. et al. Prognostic Role of Gene Mutations in Chronic Myelomonocytic Leukemia Patients Treated With Hypomethylating Agents. EBioMedicine 31, 174–181 (2018). 10.1016/j.ebiom.2018.04.018

52 Kelsey, M. M. G. Reconsidering LINE-1’s role in cancer: does LINE-1 function as a reporter detecting early cancer-associated epigenetic signatures? Evol Med Public Health 9, 78–82 (2021). 10.1093/emph/eoab004

53 Onishi-Seebacher, M. et al. Repeat to gene expression ratios in leukemic blast cells can stratify risk prediction in acute myeloid leukemia. BMC Med Genomics 14, 166 (2021). 10.1186/s12920-021-01003-z

54 Cuellar, T. L. et al. Silencing of retrotransposons by SETDB1 inhibits the interferon response in acute myeloid leukemia. The Journal of cell biology 216, 3535–3549 (2017). 10.1083/jcb.201612160

55 Gu, Z. et al. Silencing of LINE-1 retrotransposons is a selective dependency of myeloid leukemia. Nature genetics 53, 672–682 (2021). 10.1038/s41588-021-00829-8

56 Esteve, P. O. et al. Direct interaction between DNMT1 and G9a coordinates DNA and histone methylation during replication. Genes & development 20, 3089–3103 (2006). 10.1101/gad.1463706

57 Jiang, Q. et al. G9a Plays Distinct Roles in Maintaining DNA Methylation, Retrotransposon Silencing, and Chromatin Looping. Cell reports 33, 108315 (2020). 10.1016/j.celrep.2020.108315

58 Lehnertz, B. et al. The methyltransferase G9a regulates HoxA9-dependent transcription in AML. Genes & development 28, 317–327 (2014). 10.1101/gad.236794.113

59 Wingett, S. W. & Andrews, S. FastQ Screen: A tool for multi-genome mapping and quality control. F1000Res 7, 1338 (2018). 10.12688/f1000research.15931.2

60 Ewels, P., Magnusson, M., Lundin, S. & Kaller, M. MultiQC: summarize analysis results for multiple tools and samples in a single report. Bioinformatics 32, 3047–3048 (2016). 10.1093/bioinformatics/btw354

61 Langmead, B. & Salzberg, S. L. Fast gapped-read alignment with Bowtie 2. Nature methods 9, 357–359 (2012). 10.1038/nmeth.1923

62 Zhang, Y. et al. Model-based analysis of ChIP-Seq (MACS). Genome biology 9, R137 (2008). 10.1186/gb-2008-9-9-r137

63 Robinson, M. D., McCarthy, D. J. & Smyth, G. K. edgeR: a Bioconductor package for differential expression analysis of digital gene expression data. Bioinformatics 26, 139–140 (2010). 10.1093/bioinformatics/btp616

64 Heinz, S. et al. Simple combinations of lineage-determining transcription factors prime cis-regulatory elements required for macrophage and B cell identities. Molecular cell 38, 576–589 (2010). 10.1016/j.molcel.2010.05.004

65 Zhang, Y., Parmigiani, G. & Johnson, W. E. ComBat-seq: batch effect adjustment for RNA-seq count data. NAR Genom Bioinform 2, lqaa078 (2020). 10.1093/nargab/lqaa078

66 Jin, Y., Tam, O. H., Paniagua, E. & Hammell, M. TEtranscripts: a package for including transposable elements in differential expression analysis of RNA-seq datasets. Bioinformatics 31, 3593–3599 (2015). 10.1093/bioinformatics/btv422

67 Bendall, M. L. et al. Telescope: Characterization of the retrotranscriptome by accurate estimation of transposable element expression. PLoS Comput Biol 15, e1006453 (2019). 10.1371/journal.pcbi.1006453

68 Anders, S., Pyl, P. T. & Huber, W. HTSeq--a Python framework to work with high-throughput sequencing data. Bioinformatics 31, 166–169 (2015). 10.1093/bioinformatics/btu638

69 Stephens, M. False discovery rates: a new deal. Biostatistics 18, 275–294 (2017). 10.1093/biostatistics/kxw041

