## Supplementary figures and tables for "Targeting Heterochromatin Eliminates Chronic Myelomonocytic Leukemia Malignant Stem Cells Through Reactivation of Retroelements and Innate Immune pathways"

#### Supplementary materials

Supplementary Figures 1 to 11

Supplementary Tables 1 to 4

**Supplementary Fig. 1**

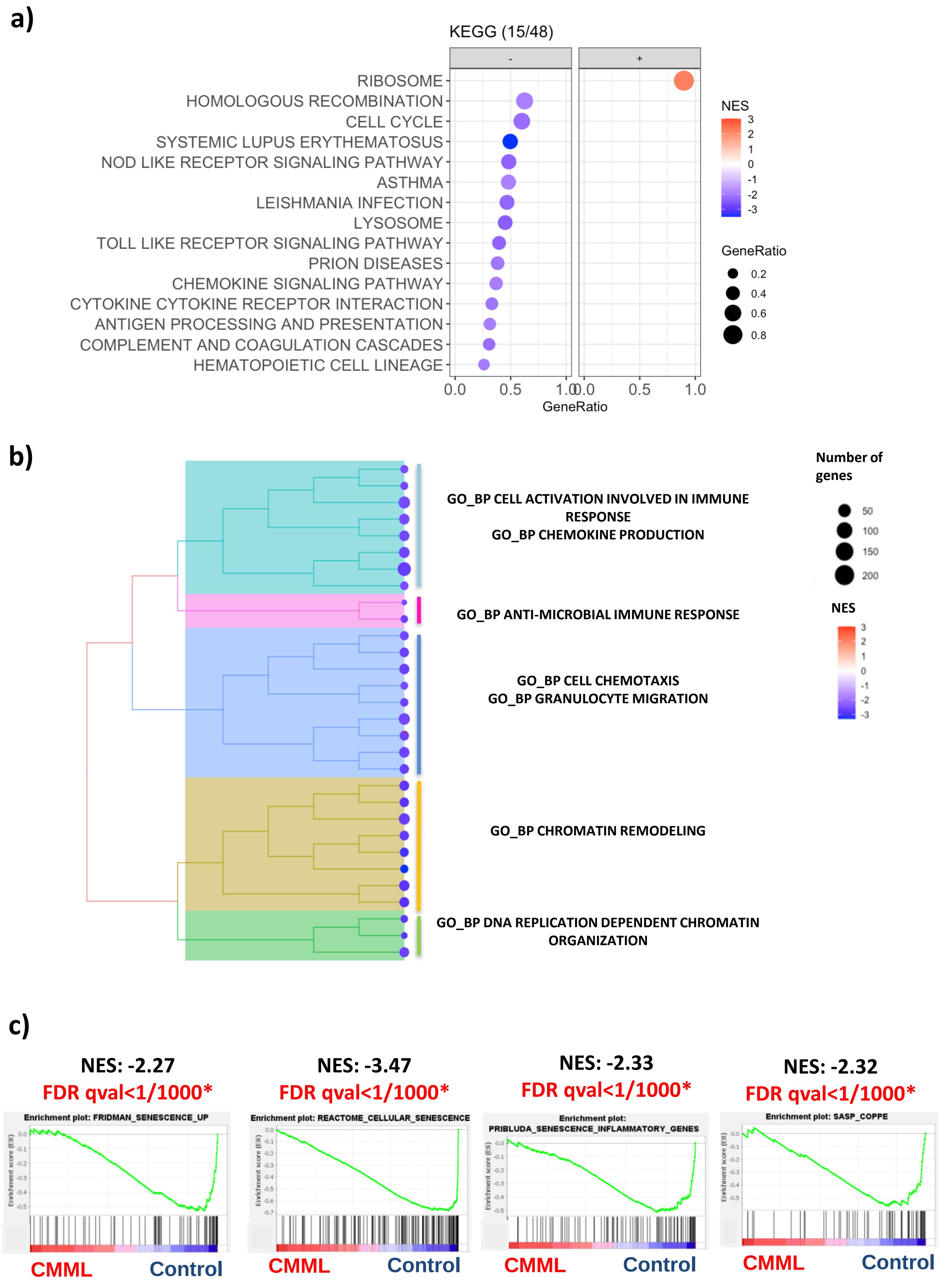

**Supplementary Fig. 1: Gene score enrichment analysis of differentially expressed genes in $CD34^+$ cells from CMML patients vs. controls**.

**a)** Dot plots showing top 15 enriched Kyoto Encyclopedia of Genes and Genomes (KEGG) pathways (FDR<0.05) upregulated (positive NES, red dots) and downregulated (negative NES, blue dots) in CMML cells compared to controls. Gene sets are ranked by their NES and gene ratio scores.

**b)** Top 30 enriched Gene Ontology Biological Processes (GO-BP) organized as a tree plot regrouping redundant pathways. NES scale and gene ratio scores are shown on the right.

**c)** Enrichment plots for published senescence and senescence-associated secretory phenotype (SASP) gene sets. FDR, false discover rate; NES, normalized enrichment score. *, FDR<0.05.

Supplementary Fig. 2

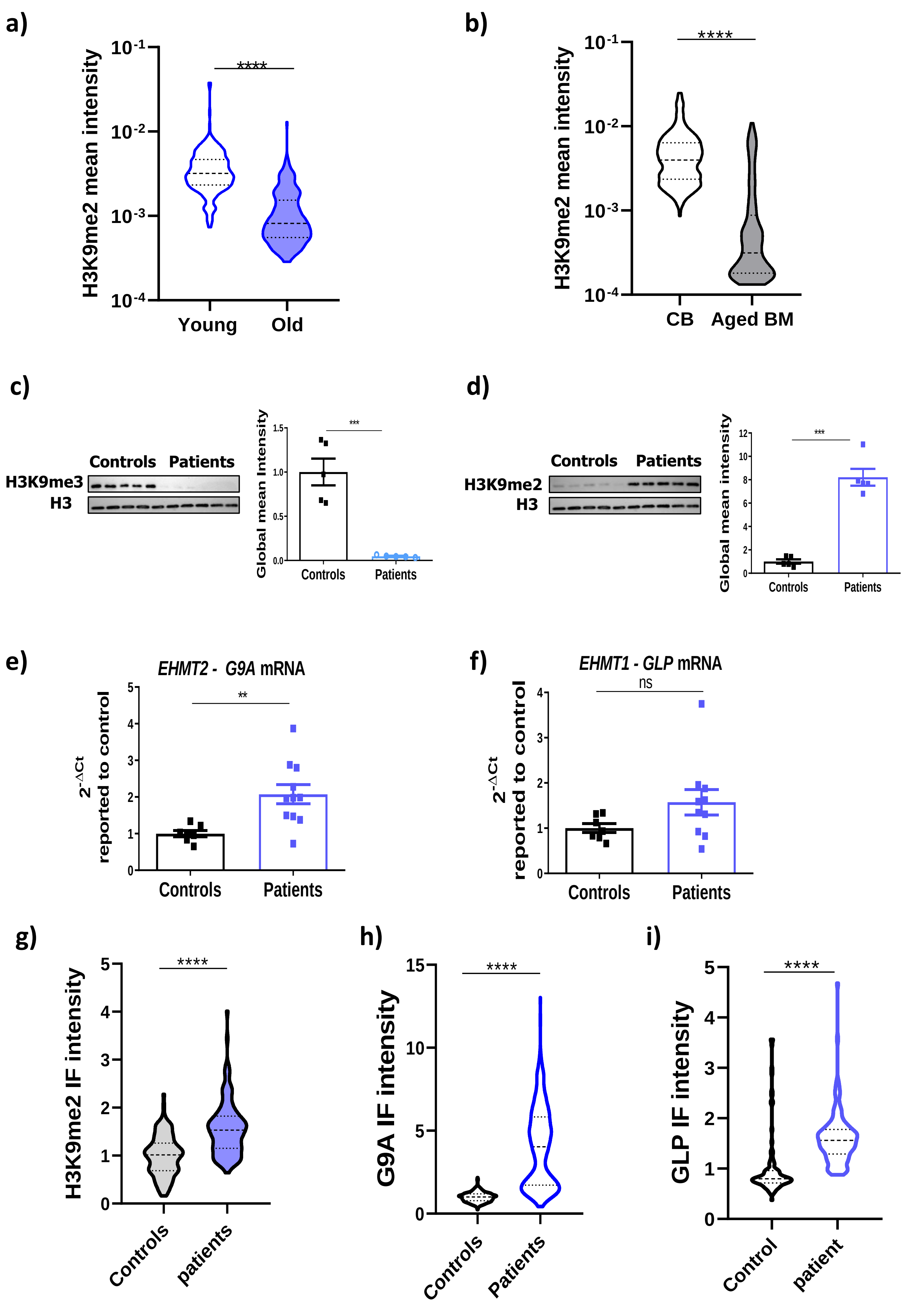

#### Supplementary Fig. 2, continued

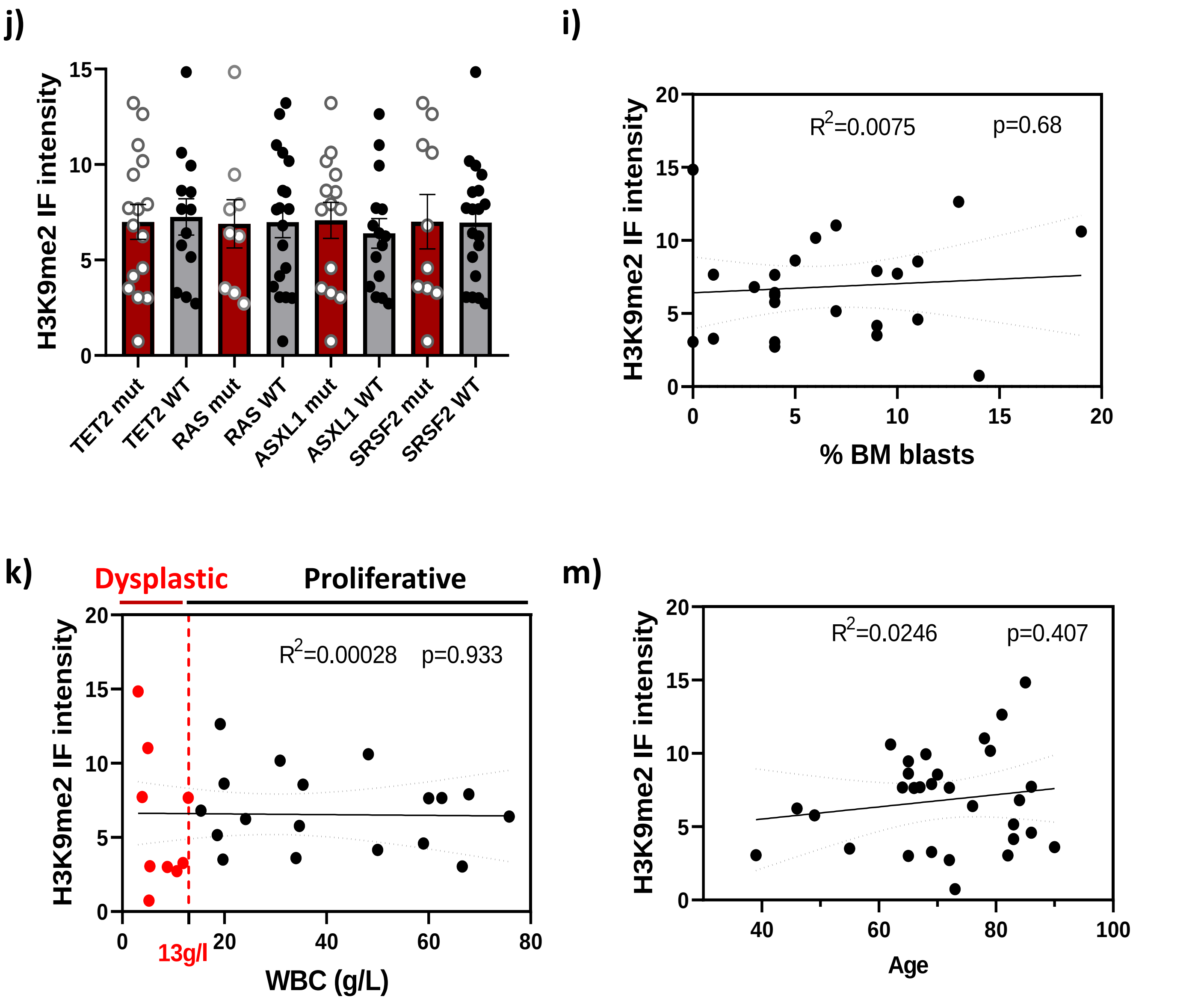


**Supplementary Fig. 2: Chromatin is remodeled in CMML HSPCs.**

**a, b)** H3K9me2 immunostaining in $Lin^{-}Sca^{+}Kit^{+}Flk2^{-}CD34^{-}$ HSCs (a) isolated from young (2 mo-old, n=3) and aged (20-24 mo-old, n=4) mice, and $CD34^{+}$ cells (b) isolated from cord blood (CB, n=4) and aged healthy donor BM (n=3). Violin plot showing the global mean intensity of each cell, from 2 independent experiments, measured by IMAGE J. 50-300 cells counted per sample. Dotted lines show median, and 25 and 75 percentiles. t-tests. ****p<0.0001.

**c, d)** Western-blot analysis of H3K9me3 (c) and H3K9me2 (d) expression relative to total H3 in $CD34^{+}$ cells from 5 controls and 5 CMML patients. t-test. **p<0.01; *** p<0.001

**e, f)** mRNA expression level measured by RT-qPCR of EHMT2, encoding G9A (e) and EHMT1, encoding GLP (f) in $CD34^{+}$ cells from controls and CMML patients. Ct values are normalized to the mean of housekeeping genes (HPRT, PPIA, Tubulin, Gus and/or RPL32) and reported to the mean of controls. Means +/- SEM from 2 independent experiments. Each dot represents cells from a single individual. t-test. **p<0.01

**g-i)** Immunostaining for H3K9me2 (g), G9A (h) and GLP (i) in $CD34^{+}CD38^{-}CD90^{+}$ from 2 (g, h) or one (i) controls and patients. Means +/- SEM IF intensity of each cell. 60-120 cells analyzed per sample. Dotted lines: median and 25th and 75th percentiles. t-test. ****p<0.0001

**J-m)** Correlation of H3K9me2 mean IF intensity levels with the presence of TET2, RAS, ASXL1 and SRSF2 mutations (j), white blood cell (WBC) counts (k), the percent pf blasts in the bone marrow (BM) (l) and age (m). Dysplastic forms: <13g/l WBC; Proliferative forms >13g/l WBC counts.

#### Supplementary Fig. 3

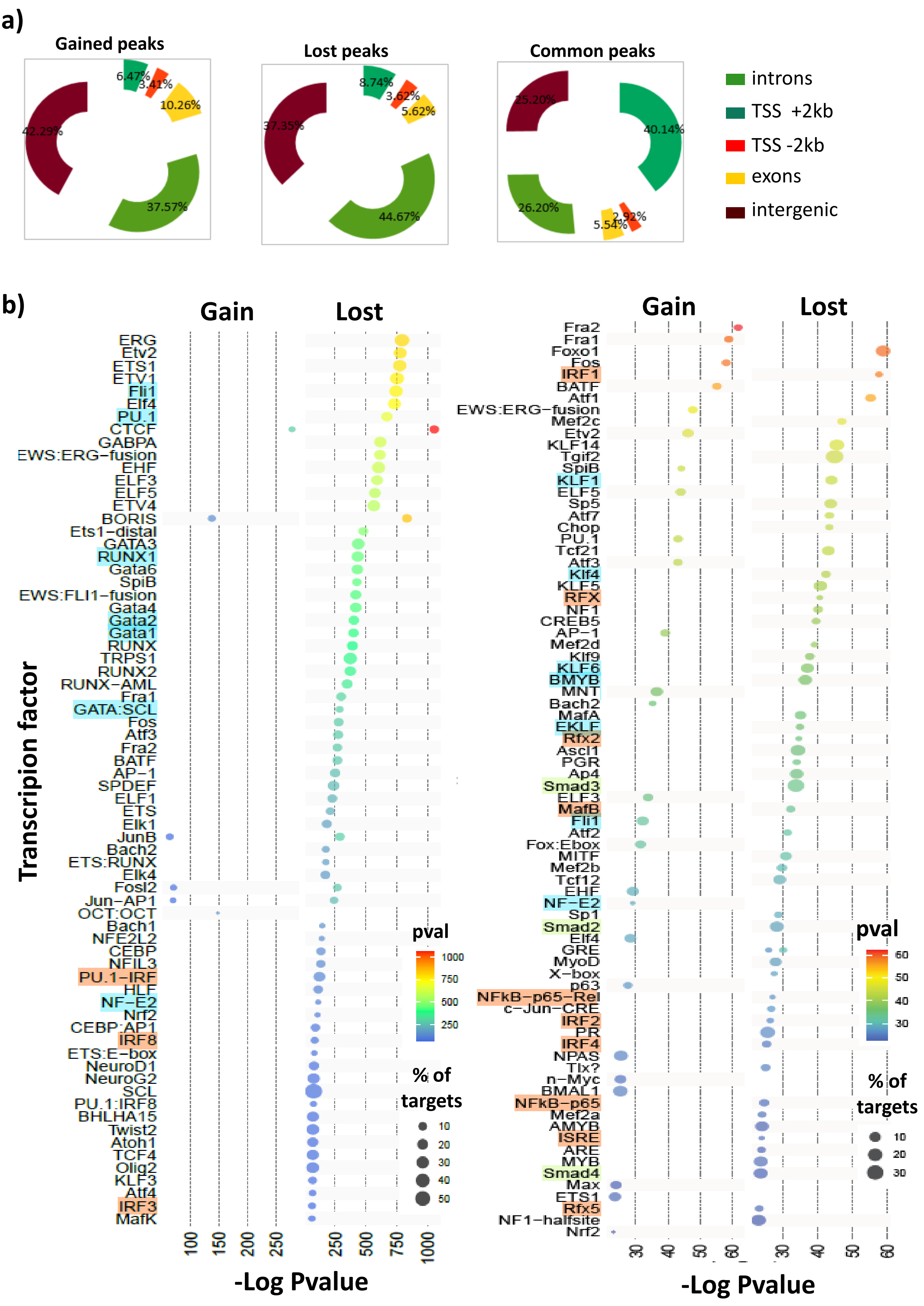

**c)**

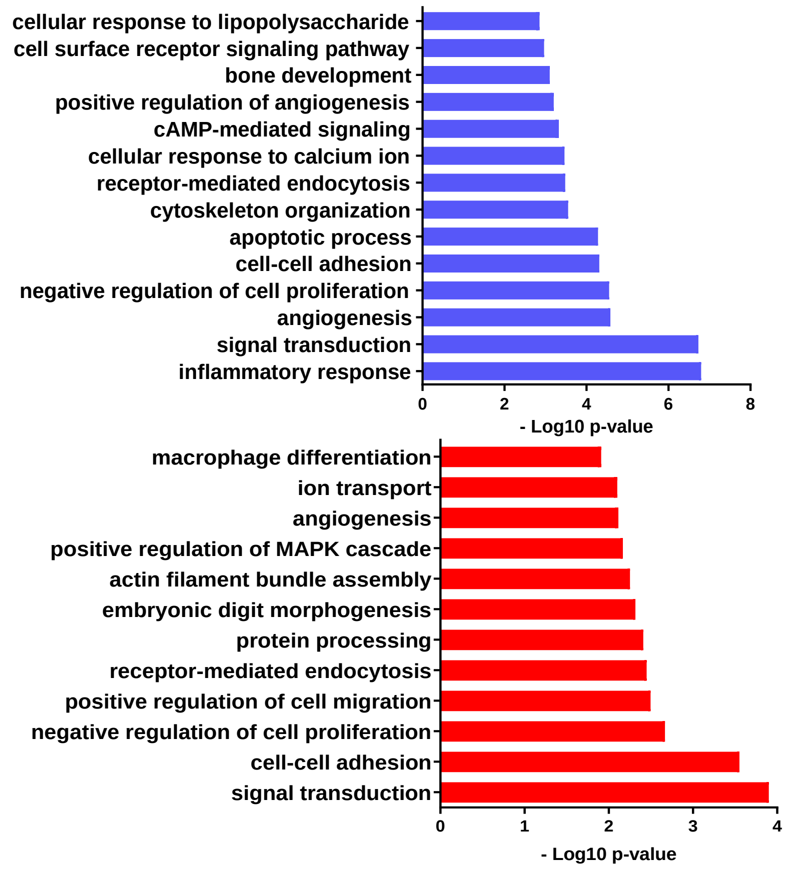


**Supplementary Fig. 3: Chromatin is remodeled in CMML HSPCs.**

**a)** Genomic repartition of ATAC-seq peaks uniquely found in patients (gained chromatin accessibility) or in controls (lost chromatin accessibility) or in common peaks.

**b)** Transcription factor binding site enrichment analyzed by HOMER in ATAC-seq peaks gaining or losing accessibility in CMML patients as compared to controls. Factors are ranked by pvalue (see Supplementary Data 4). Left: p<$10^{-27}$; right: >$10^{-27}$ – $10^{-10}$. The color of the dots represents the pvalue; the size of the dots represents the % of binding site targets. Transcription factors involved in hematopoietic differentiation (blue), IFN/NF-κB signaling pathways (orange) and TGF-β signaling (yellow) are highlighted.

**c)** KEGG gene set analysis of deregulated genes associated with CMML ATAC peaks that are lost (upper panel, blue) or gained (lower panel, red) in CMML patient cells.

### Supplementary Fig. 4

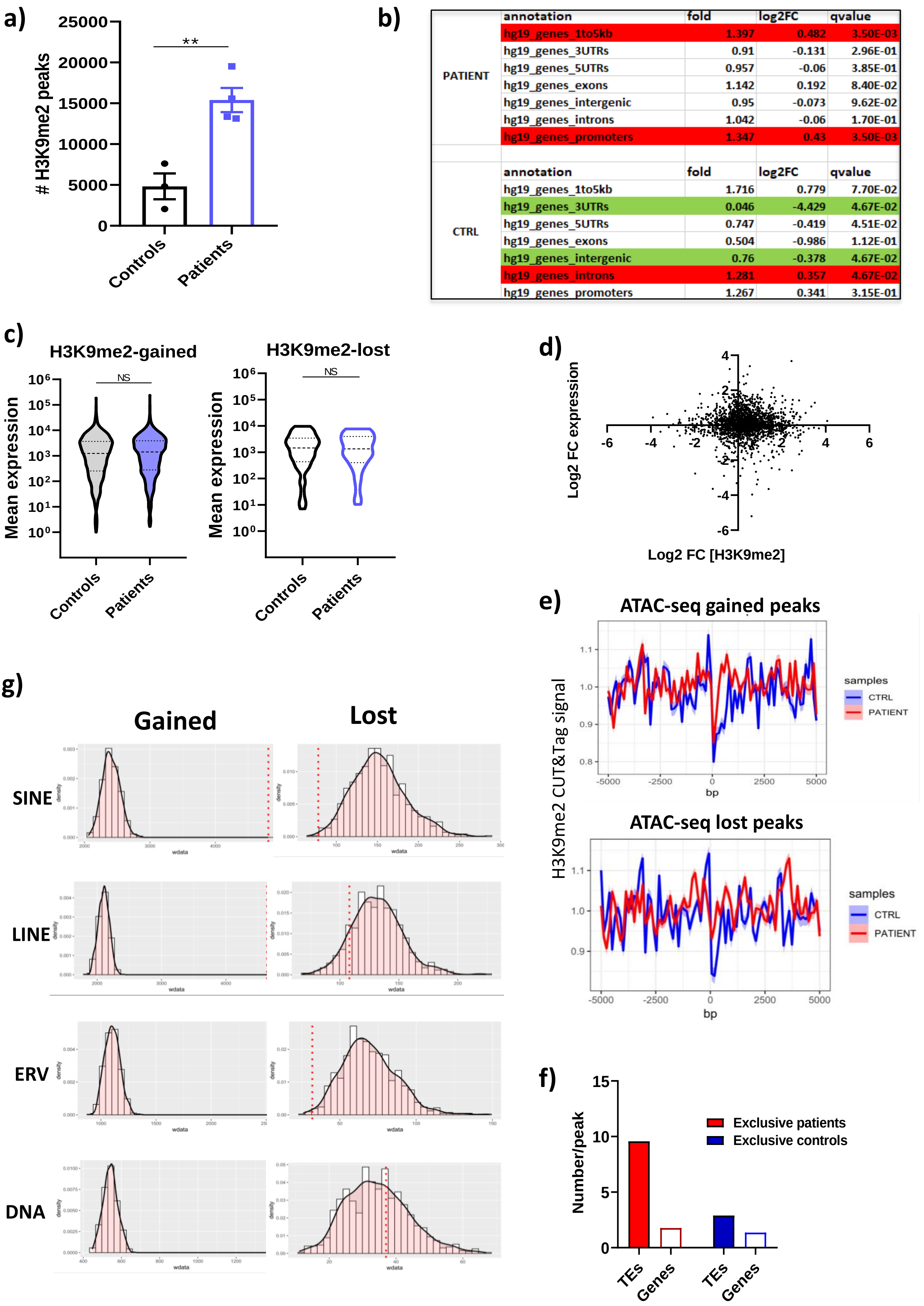


| | annotation | fold | log2FC | qvalue |
|---|---|---|---|---|
| PATIENT | hg19_genes_1to5kb | 1.397 | 0.482 | 3.50E-03 |
| | hg19_genes_3UTRs | 0.91 | -0.131 | 2.96E-01 |
| | hg19_genes_5UTRs | 0.957 | -0.06 | 3.85E-01 |
| | hg19_genes_exons | 1.142 | 0.192 | 8.40E-02 |
| | hg19_genes_intergenic | 0.95 | -0.073 | 9.62E-02 |
| | hg19_genes_introns | 1.042 | -0.06 | 1.70E-01 |
| | hg19_genes_promoters | 1.347 | 0.43 | 3.50E-03 |
| | annotation | fold | log2FC | qvalue |
| CTRL | hg19_genes_1to5kb | 1.716 | 0.779 | 7.70E-02 |
| | hg19_genes_3UTRs | 0.046 | -4.429 | 4.67E-02 |
| | hg19_genes_5UTRs | 0.747 | -0.419 | 4.51E-02 |
| | hg19_genes_exons | 0.504 | -0.986 | 1.12E-01 |
| | hg19_genes_intergenic | 0.76 | -0.378 | 4.67E-02 |
| | hg19_genes_introns | 1.281 | 0.357 | 4.67E-02 |
| | hg19_genes_promoters | 1.267 | 0.341 | 3.15E-01 |

**Supplementary Fig. 4: H3K9me2 CUT&Tag analysis**

**a)** number of H3K9me2-bound peaks detected in the cells from the 3 controls and the 4 patients. Means +/- SEM; t-test. **$p<0.01$.

**b)** Genomic Association Test showing the enrichment of H3K9me2 in genomic annotations in patients and controls.

**c)** Mean expression levels of genes linked to H3K9me2 gained and lost peaks in CMML patients.

**d)** Correlation plot representing the $\log_2$ fold-change [FC] in H3K9me2 concentration at gene promoters vs. $\log_2$ fold-change in gene expression between patients and controls.

**e)** Normalized read distribution profiles of H3K9me2 CUT&Tag in patients and controls spanning gained (upper panel) or lost (lower panel) ATACseq peaks, +/- 5 kb from the center of the peak (0).

**f)** Numbers of TEs (filled bars) or genes (open bars) in H3K9me2-bound peaks that are found exclusively in the 4 patients (red) and 3 controls (blue).

**g)** Permutation test comparing of the number of patient (gained) and control (lost) specific H3K9me2-bound peaks overlapping with TEs specific and in 1000 lists of identical numbers of random peaks. Black curve: Distribution of the number of peaks overlapping TEs found in random peaks. Red vertical dotted line: number of peaks overlapping TEs in the differential peaks.

#### Supplementary Fig. 5

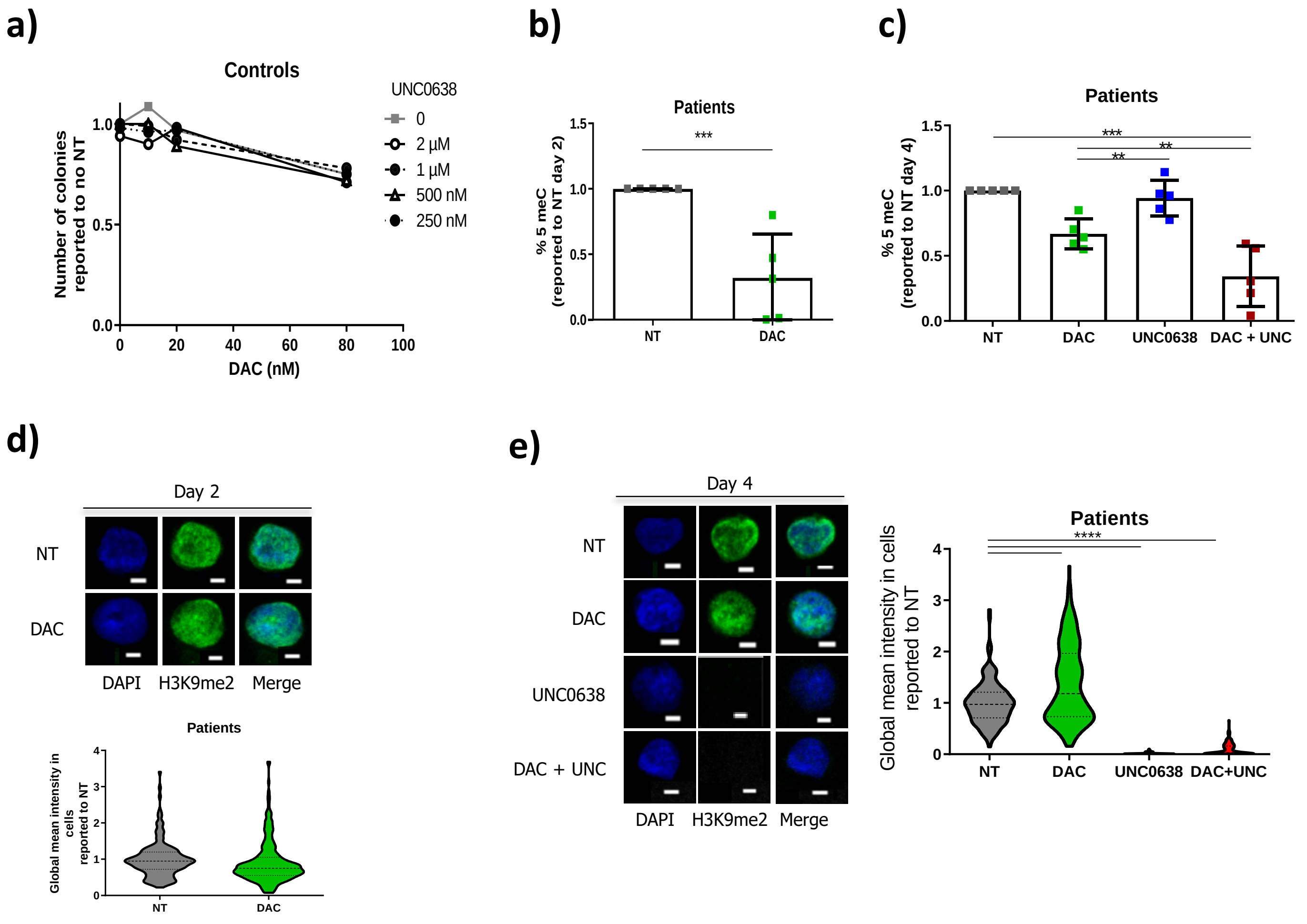


**Supplementary Fig. 5: Specificity of HMAs and G9A/GLP inhibitors.**

**a)** Dose-response effect of DAC (X-axis) and UNC0638 (dark/grey scale) on CD34+ colonies formed by cells from one control.

**b, c)** Measurement of DNA methylation level (% 5meC) after 2 (b) and 4 (c) days of liquid culture of CMML $CD34^+$ cells in the presence or absence of DAC (10 nM) and UNC0638 (1µM), as indicated. Means +/- SEM, n=5 patients. Results are normalized to the percent of 5meC in the NT condition.

**d, e)** Representative images and quantification of H3K9me2 IF mean intensity from CMML patient (n=3) $CD34^+$ cells after 2 (d) and 4 (e) days of liquid culture reported to the mean intensity of non-treated cells. Bars, 3 µm. Images were quantified using ImageJ software.

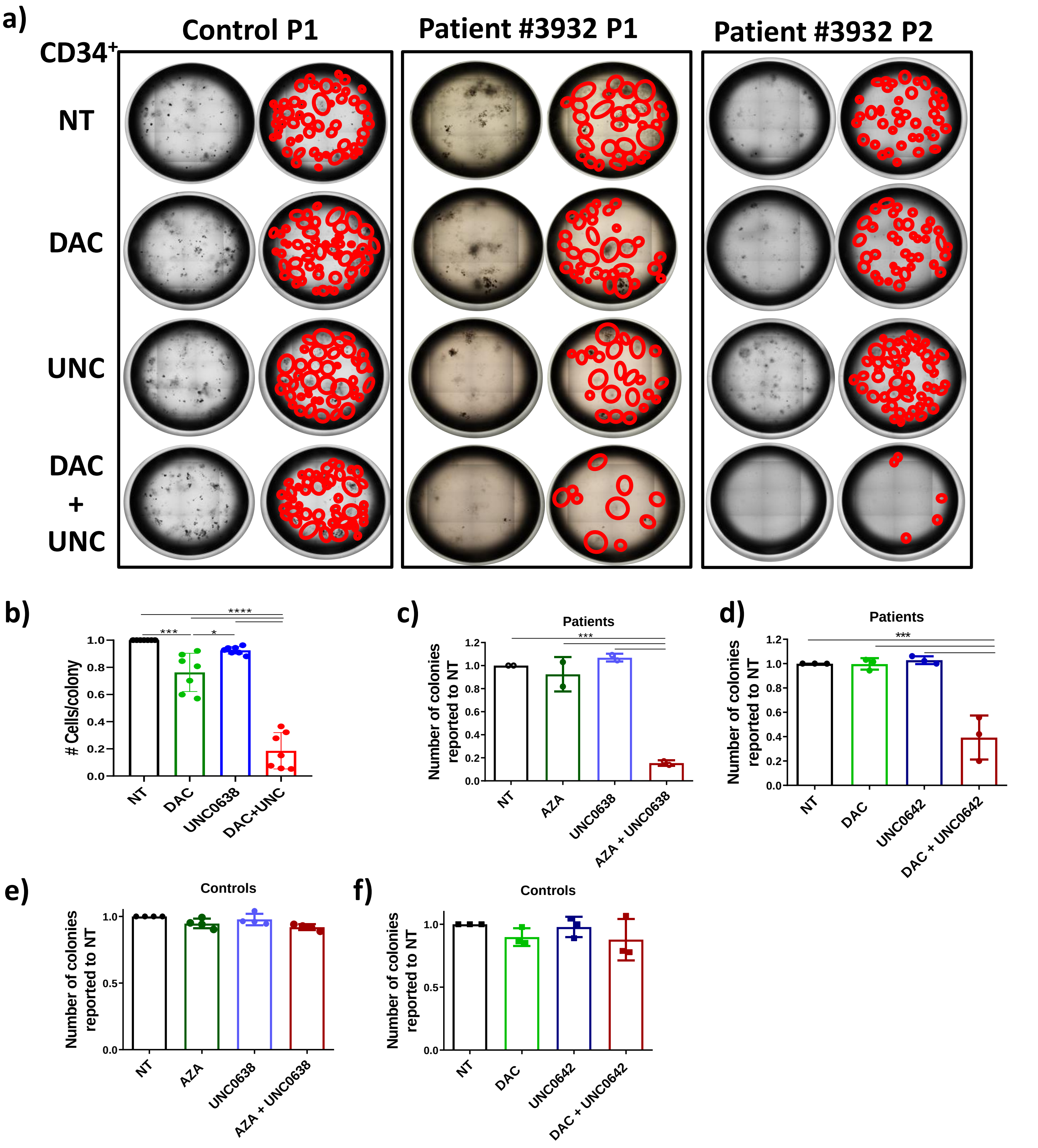


**Supplementary Fig. 6: The combination of HMAs and G9A/GLP inhibitors induce a specific loss of CMML CD34$^+$ cell clonogenicity.**

**a)** Raw images of colonies formed from a control and a patient in the presence or absence of DAC, UNC0638 or both at passage 1 (P1) or 2 (P2), as indicated. The first columns of each panel show the raw images of the whole plate. In the second columns, each colony counted was circled in red.

**b)** Number of cells per colony formed by CD34$^+$ cells from CMML patients treated with DAC, UNC0638 or both.

**c-f)** Number of colonies formed from patient (c, d) or control (e, f) CD34$^+$ cells after culture with 5-AZA-cytidine (AZA, 500 nM) in the presence of absence of G9A/GLP inhibitor UNC0638 (1 µM) (left panel) or the combination of DAC (10 nM) with the G9A/GLP inhibitor UNC0642 (2 nM) (right panel). One dot represents a single individual. Means +/- SEM. One-way ANOVA Bonferroni's Multiple Comparison *p<0.05; ***p<0.001.

#### Supplementary Fig. 7

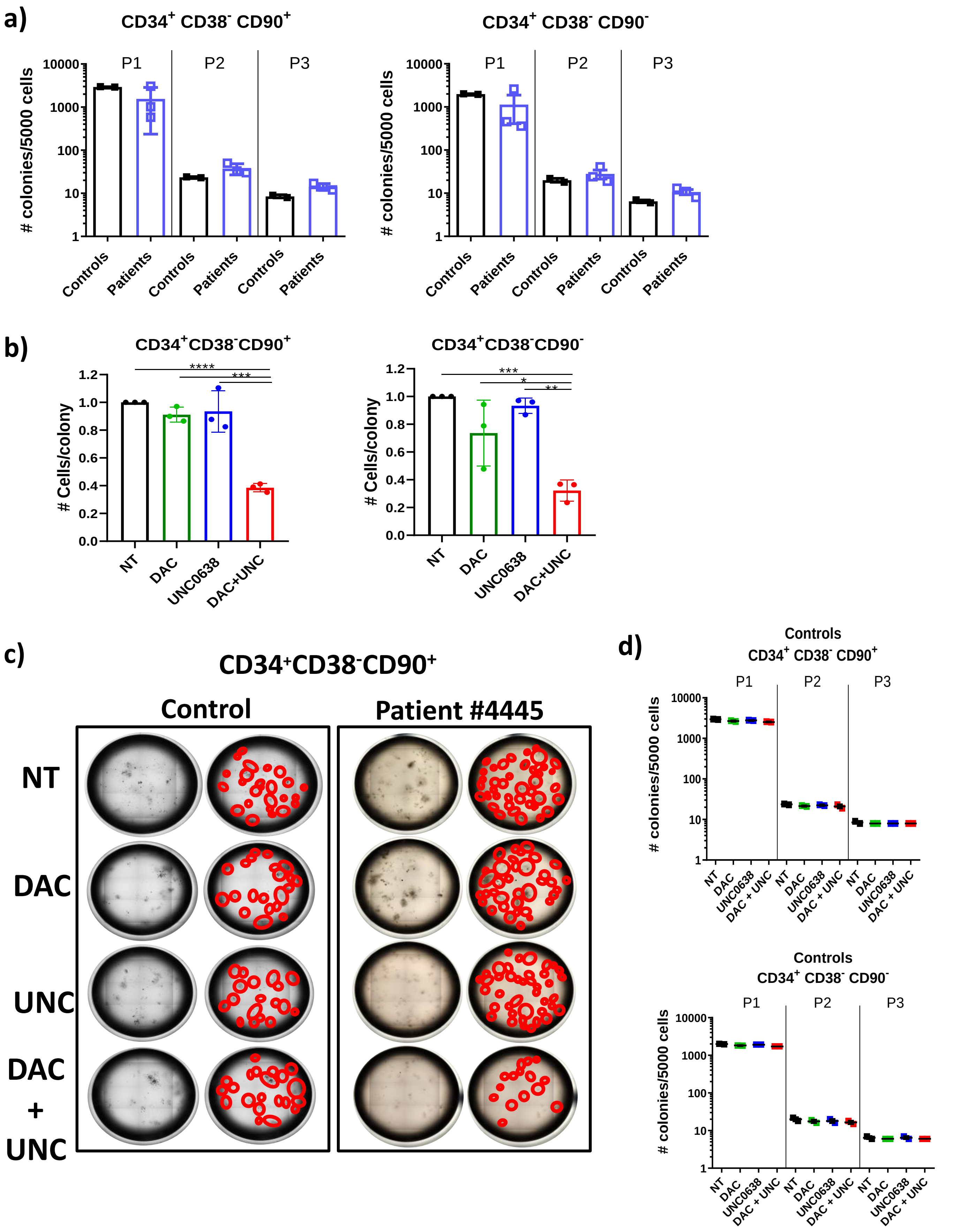

**Supplementary Fig. 7: The HMA and G9A/GLP inhibitor combination reduces CMML mutated stem cells clonogenicity**.

**a)** Number of colonies formed at repeated passages from stem cells ($CD34^{+}CD38^{-}CD90^{+}$, left panel) or progenitors ($CD34^{+}CD38^{-}CD90^{-}$, right panel) from CMML patients and controls before treatment. One dot represents a single individual

**b)** Number of cells per colony formed by CMML patient stem cells (left panel) or progenitors (right panel) treated with DAC, UNC0638 or both, or left untreated. Results are reported to the mean number of cell/colony found in the untreated samples. Means +/- SEM. One dot represents a single individual One-way ANOVA Bonferroni's Multiple Comparison.

**c)** Raw images of colonies formed from CD34+CD38-CD90+ from one control and one patient in the presence or absence of DAC, UNC0638 at passage 1. The first column of each panel show the raw images of the whole plates. In the second columns, each colony counted was circled in red.

**d)** Number of colonies formed at passages 1 to 3 by stem cells and progenitors from controls, in the presence or absence of treatment. Means +/- SEM. One dot represents a single individual One-way ANOVA Bonferroni's Multiple Comparison.

### Supplementary Fig. 8

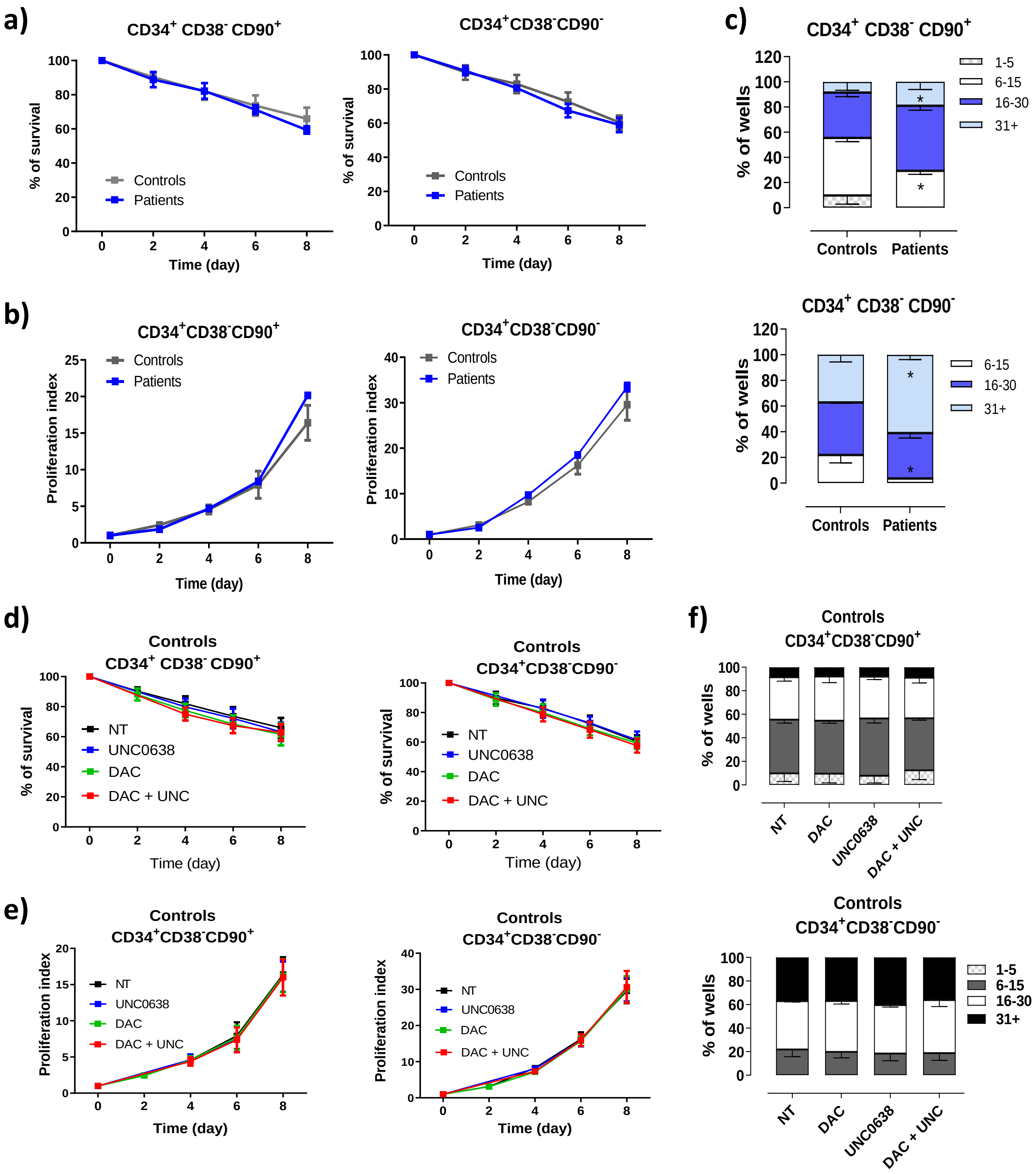


**Supplementary Fig. 8: The HMA and G9A/GLP inhibitor combination reduces the survival and the proliferation CMML mutated stem cells in single cell liquid culture**.

**a-b)** Single cell liquid cultures comparing the survival (a), the proliferation (b) and the cell of stem cells ($CD34^{+}CD38^{-}CD90^{+}$) and progenitors ($CD34^{+}CD38^{-}CD90^{-}$) from patients (n=3, blue) or control cells (n=3, grey). Means +/- SEM.

**c)** Cell number repartition per well after 14days of culture of control and patient stem cells and progenitors as indicated. Two-way ANOVA Bonferroni's Multiple Comparison. *p<0.05;

**d-f)** Viability, (d) proliferation index, (e) and cell number repartition per well (f) of control cells treated with either DAC, UNC0638 or both, or left non-treated (n-=3). Means +/- SEM.

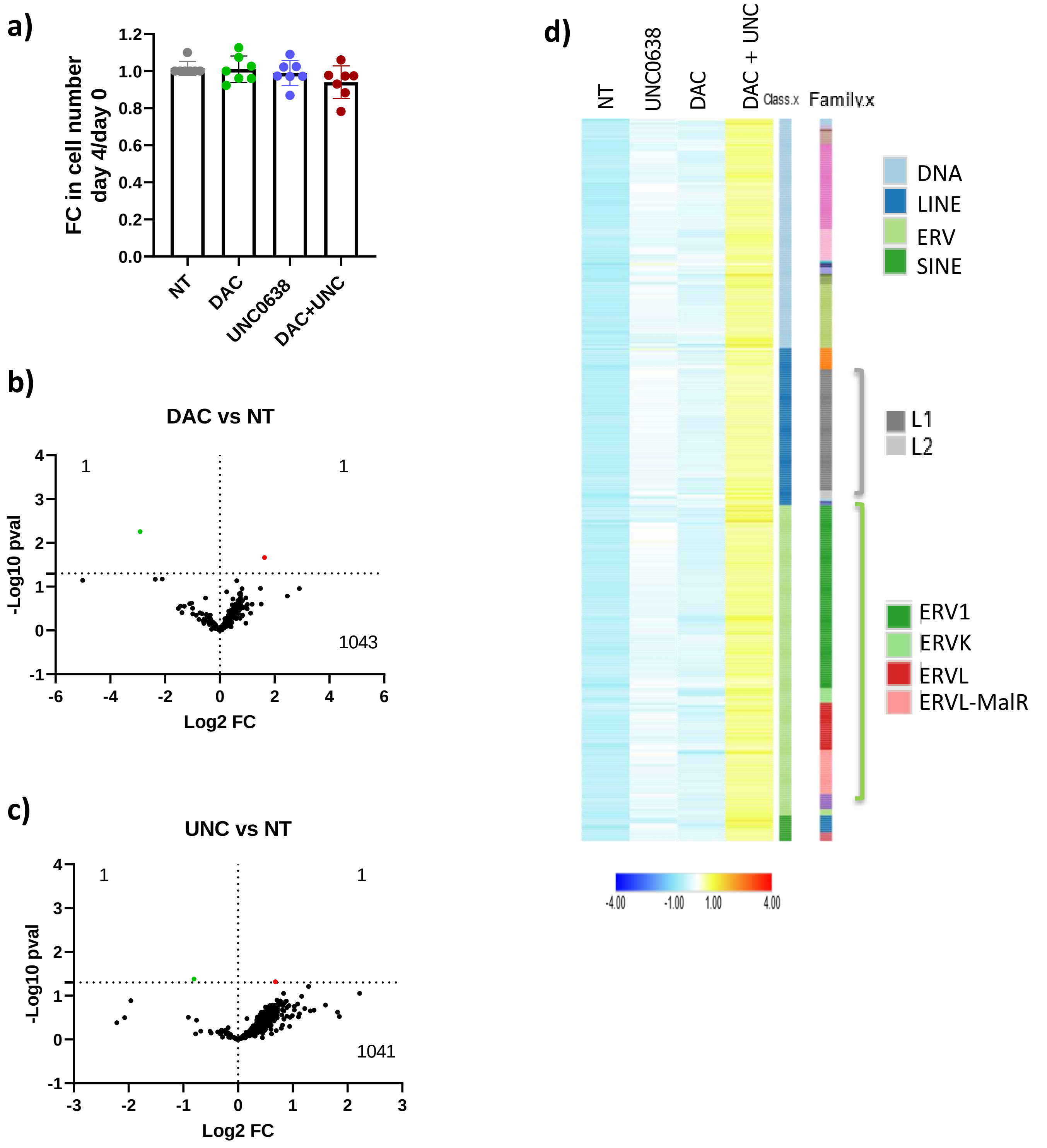


**Supplementary Fig. 9: The HMA and G9A/GLP inhibitor combination increases TE expression in CMML CD34⁺ HSPCs.**

**a)** Number of cells remaining in the cultures at day 4 in the presence of the various inhibitors. Fold-change in cell numbers reported to the number of cells seeded at day 0.

**b, c)** Volcano plots showing differential TE expression in cells treated with DAC (b) or UNC0638 (c), compared to NT. Red and green dots: upregulated and downregulated TEs, respectively.

**d)** Heatmap showing the median expression of DAC+UNC0638 vs. NT differentially expressed TEs in each of the treatment conditions, clustered on both TE class and TE family.

### Supplementary Fig. 10

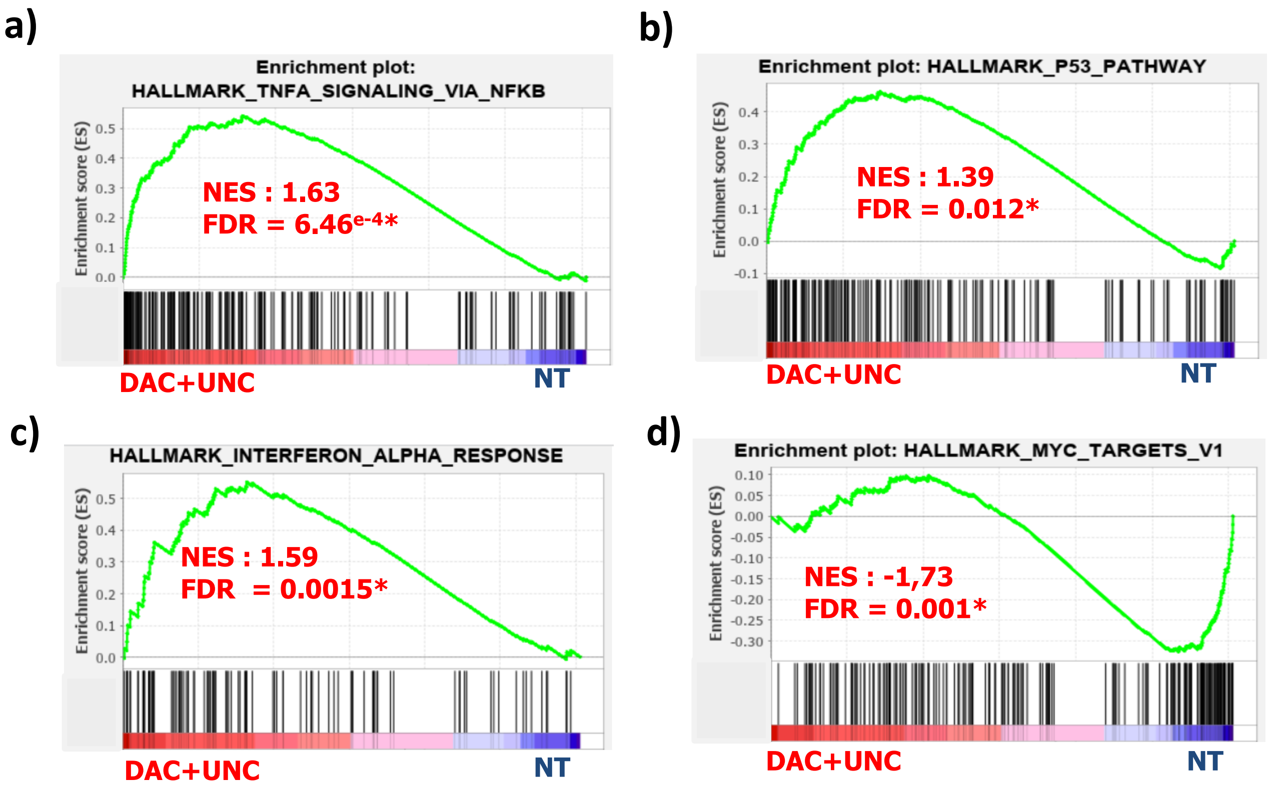


**Supplementary Fig. 10: The DAC/UNC0638 combination reactivates viral and inflammatory pathways in CMML cells.**

GSEA enrichment plots for selected pathways upregulated **(a-c)** or downregulated **(d)** in CMML cells after treatment with DAC+UNC0638. FDR, false discover rate; NES, normalized enrichment score. *, FDR<0.05.

### Supplementary Fig. 11

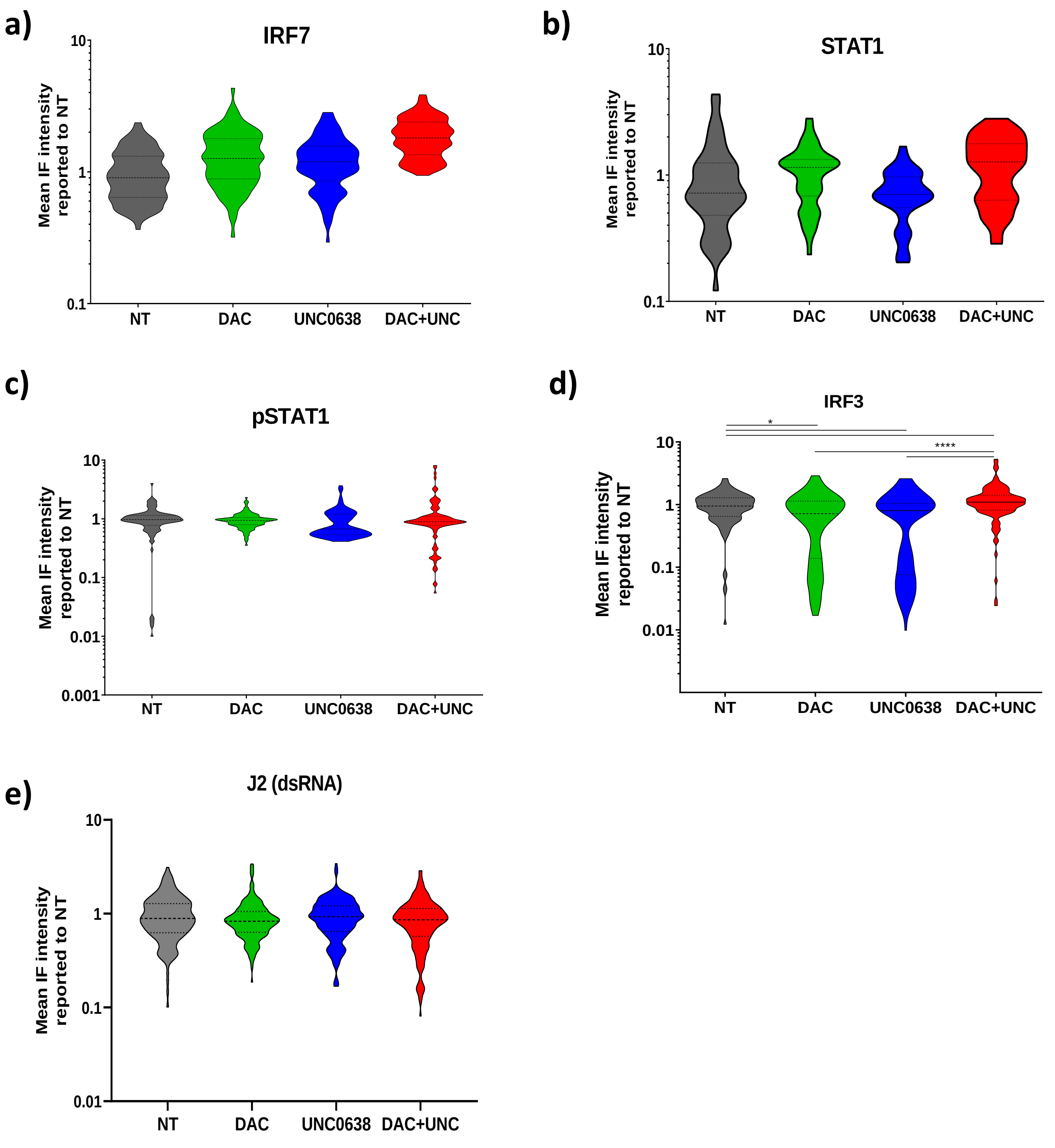


**Supplementary Fig. 11: The DAC/UNC0638 combination does not trigger IFN signaling in control cells**

**a-e)** Immunostaining with antibodies against IRF7 (a), STAT1 (b), pSTAT1 (c), IRF3 (d) and dsRNAs (e) in $CD34^+$ cells from one (a, b) or two (c, d, e) healthy donors after treatment with DAC +/- UNCO638, or untreated. Violin plots showing the global mean intensity of each cell, reported to the NT condition. 45-130 cells analyzed per sample. Dotted lines: median and 25th and 75th percentiles. One-way ANOVA Bonferroni's Multiple comparison. *p<0.05; ***p<0.001.

**Supplementary Table 1: CMML patient and healthy donor cohorts**

| | Cohort | |
|---|---|---|
| | **Healthy controls** | **Patients** |
| Number<br>Males, N (%) | 39<br>16 (42%) | 100<br>54 (54%) |
| Age, median [range] | 68 [45-91] | 72 [40-90] |
| CMML-1 (%)<br>CMML-2 (%) | | 78%<br>22% |
| Dysplastic (%)<br>Proliferative (%) | | 42%<br>58% |
| Leukocytosis G/L; median [range] | | 15.8 [3.07-157.2] |
| Hemoglobin g/dL; median [range] | | 11.4 [4.5-16.6] |
| Platelets, G/L; median[range] | | 119 [11.6-986] |
| Neutrophils, G/L; median [range] | | 7.9 [0.6-95.9] |
| Monocytes, G/L; median G/L [range] | | 3.2 [0.65-53.2] |
| Monocytes, %; median [range] | | 20.4 [5-57.7] |
| Bone marrow blasts, median % [range] | | 5 \|0-19] |
| Next generation sequencing analysis ; N (%) | | 100 |
| ASXL1 | | 39 |
| BRCC3 | | 1 |
| CBL | | 15 |
| CSF3R | | 3 |
| CTCF | | 1 |
| CUX1 | | 6 |
| DNMT3A | | 12 |
| ETNK1 | | 1 |
| ETV6 | | 2 |
| EZH2 | | 7 |
| IDH1 | | 5 |
| IDH2 | | 4 |
| KDM6A | | 2 |
| KIT | | 1 |
| KRAS | | 13 |
| NRAS | | 14 |
| PHF6 | | 4 |
| PTPN11 | | 1 |
| RIT1 | | 1 |
| RUNX1 | | 15 |
| SF3B1 | | 7 |
| SRSF2 | | 41 |
| TET2 | | 65 |
| TP53 | | 2 |
| U2AF35 | | 3 |
| U2AZF1 | | 1 |
| ZRSR2 | | 3 |

**Supplementary Table 2: Mutations and variant allele frequencies in 5 patients**

| Patient #3440 | Patient #3480 | Patient #3459 | Patient #3562 | Patient #3564 |
|---|---|---|---|---|
| **RUNX1** NM_001001890.2 exon 2 c.341C>T : p.Ser114Leu (61%) | **TET2** NM_001127208.2 exon 9 c.4139A>G : p.His1380Arg (37%) | **TET2** NM_001127208.2 exon 9 c.4140T>A : p.His1380Gln (44%) | **KRAS** c.G35A: p.G12D (30%) | **TET2** NM_001127208.2 exon:6 c.3686T>C p.Leu1229Pro (66%) |
| **TET2** NM_001127208.2 exon 8 c.4012A>T : p.Lys1338*, nonsense mutation, (54%) | **DNMT3A** NM_022552.4 exon 8 c.1014+1delG : splice_donor_variant (34%) | | | **SRSF2** NM_001195427/1 Exon 1 c.284C>G p.Pro95Arg (37%) |

**Supplementary Table 3: Primers used in this study**

Primers for VAF analysis

| Gene | Sense | Anti-sense |
|---|---|---|
| RUNX1-S114L (Patient #3340) | GTCCTTTGACTGGTGTTTAGG | GCATAGTTTTGACAGATAACG |
| TET2-K1338X (Patient #3340) | CTAACTGATAGTCTCTTTTACATAGAG | GTAACAAGTAAGTTGTTACAATTG C |
| TET2-H1380R (Patient #3380) | CAGATTGAATATGAACACAGAGC | CAGACTCAACAGCTGCTAAGC |
| DNMT3A c.1014+1 delG (Patient #3380) | GCATTGTGTCTTGGTGGATG | CATCACCCCAATTCCAGACTG |
| TET2-L1229P (Patient #3564) | GTGGTTCGCAGAAGCAGCAG | CATTCAAGGCACACCGGCG |
| SRSF2-P95R (Patient #3564) | TGTAAAACGACGGCCAGTGGCCGCCAC TCAGAGCTA | GGAAACAGCTATGACCATGCGCGG ACCTTTGTGAGGT |
| TET2-H1380Q (Patient #3459) | CAGATTGAATATGAACACAGAGC | CAGACTCAACAGCTGCTAAGC |
| KRAS-G12D (Patient #3562) | GTGTGCATGTTCTAATATAGTC | CTATTGTTGGATCATATTCGTCC |

Primers for RT-QPCR analysis

| Gene | Sense | Anti-sense |
|---|---|---|
| EHMT2 | GGCCCCCTGAAATACAGCAT | TCGTCAGGGTCACTTCACCT |
| EHMT1 | GTGTGAAAACCGAGCTGCTG | TCCTGCCTGTTTCTCTGCTG |
| ERVL | ATATCCTGCCTGGATGGGGT | GAGCTTCTTAGTCCTCCTGTGT |
| ERVLB4 | CAGTCAGCCTCTTTCCCCAG | TGTTGCTGAGCCCATGCATA |
| MER4D0 | TCCGGAGAGTCTGACACCTT | CCAGGTAGCAGGCTTCAGAG |
| ERVH | CCCTGTCCTCCTGCTCTTTG | TGGTGAGATGTTCCTTGGGC |
| ALUYH9 | CGCCTGTAATCCCAGCACTT | GCCAGGATGGTCTCGATCTC |
| IFNB1 | AAACTCATGAGCAGTCTGCA | AGGAGATCTTCAGTTTCGGAGG |
| IRF7 | CGAGACGAAACTTCCCGTCC | GCTGCCTCGGTATGGATCTC |
| IRF9 | GTCCAGCTGTCTGGAAGACTC | GGTACTTTCTGAGTCCCTGGC |
| GUS | GAAAATATGTGGTTGGAGAGCTCATT | CCGAGTGAAGATCCCCTTTTTA |
| HPRT | GGACAGGACTGAACGTCTTGC | CTTGAGCACACAGAGGGCTACA |
| TUBULINE | TCCAGATTGGCAATGCCTG | GGCCATCGGGCTGGAT |
| PPIA | GTCGACGGCGAGCCC | TCTTTGGGACCTTGTCTGCAA |
| RPL32 | TGTCCTGAATGTGGTCACCTG | CTGCAGTCTCCTTGCACACCT |

**Supplementary Table 4: Primary and secondary antibodies used in immunofluorescence and western blots**

| Specificity | **Référence** | Host | Dilution | Fluorochrome |
|---|---|---|---|---|
| G9A | GTX129153 *(GeneTex)* | Rabbit | 1/200 | |
| GLP | HPA060022 *(Altas antibody)* | Rabbit | 1/200 | |
| IRF3 | TA311894 *(Origen)* | Rabbit | 1/100 | |
| IRF7 | HPA052757 *(Sigma-Aldrich)* | Rabbit | 1/200 | |
| DsRNA | MABE1134-J2 *(Millipore)* | Rabbit | 1/150 | |
| H3K9me2 | 39239 *(Active Motif)* | Rabbit | 1/500 | |
| H3K9me3 | 8898 *(abcam)* | Rabbit | 1/1000 | |
| H3 | 18521 *(abcam)* | Rabbit | 1/5000 | |
| STAT1 | 14994 *(Cell Signaling)* | Rabbit | 1/400 | |
| pSTAT1 (Tyr701) | 9167 *(Cell Signaling)* | Rabbit | 1/400 | |
| Rabbit | A11008 *(ThermoFisher Scientific)* | Goat | 1/600 | AlexaFluor 488 |
| Mouse | A21425 *(ThermoFisher Scientific)* | Goat | 1/600 | AlexaFluor 555 |
